# A transgenic zebrafish for spatiotemporal optogenetic activation of FGF/ERK signaling

**DOI:** 10.64898/2026.05.12.724650

**Authors:** William K. Anderson, Leanne E. Iannucci, Ninet Sinaii, Julia Porcino, Katherine W. Rogers

## Abstract

Dynamic FGF/ERK signaling plays key roles in development, regeneration, and disease. We recently developed a zebrafish-optimized optogenetic tool, bOpto-FGF, that enables reversible activation of FGF/ERK signaling in response to blue light (∼455 nm) by fusing a zebrafish receptor tyrosine kinase domain to the blue light-dimerizing LOV domain. Previously, this tool was introduced into zebrafish embryos by mRNA injection. Here, we develop *Tg(ubi:bOpto-FGF)*, a novel transgenic zebrafish ubiquitously expressing bOpto-FGF, to streamline experimental workflows. We demonstrate robust blue light-mediated activation of FGF/ERK signaling and FGF target gene expression in early gastrulation-stage *Tg(ubi:bOpto-FGF)* embryos. Light-mediated signaling activation at this stage is more spatially uniform in transgenics compared to mRNA-injected embryos, and ectopic signaling in response to continuous light exposure is activated within 3 minutes and maintained for at least 75 minutes. Further, *Tg(ubi:bOpto-FGF)* heterozygotes are light-responsive from late blastula stages through at least 24 hours post-fertilization. Finally, we demonstrate light-mediated spatially localized FGF/ERK signaling activation within 1 minute in the head, trunk, and tail at 1 day post-fertilization. This transgenic line provides a powerful and convenient new strategy for spatiotemporal manipulation of FGF/ERK signaling dynamics in the vertebrate zebrafish model.

**Summary Statement:** Novel transgenic zebrafish provides precise optogenetic control of a key developmental signaling pathway *in vivo*

## INTRODUCTION

Fibroblast Growth Factor (FGF) ligand and its effector Extracellular Signal-Regulated Kinase (ERK) regulate many developmental processes across the animal kingdom. In the popular vertebrate model zebrafish (*Danio rerio*), FGF/ERK signaling dynamics play an important role during development of the germ layers, heart, brain, spinal cord, ear, skin, and sensory organs. For example, an FGF signaling gradient in the early embryo contributes to germ layer patterning by specifying mesodermal cell fates (Bökel and Brand, 2013; Jones and Mullins, 2022). At this stage high levels of FGF also promote specification of ventricular cells important for heart development, and at later stages FGF signaling reinforces ventricular cardiomyocyte identity (Yao et al., 2021). *fgf8* ligand is expressed dynamically in the developing zebrafish brain (Moens and Prince, 2002) and contributes to the patterning activities of the midbrain-hindbrain boundary (Gibbs et al., 2017). In the trunk, tissues that will contribute to the spinal cord (somites and notochord) experience ERK signaling oscillations required for normal patterning (Adhyapok et al., 2025; McDaniel et al., 2024). Ear cell precursors are induced and expanded by spatiotemporally dynamic expression of multiple FGF ligands, and distinct auditory and vestibular structures appear to be specified by low and high FGF signaling levels, respectively (Riley, 2021). Expansion of the skin during development is influenced by ERK pulses (Ramkumar et al., 2025), and the lateral line (a flow-detecting sensory system) is formed by a group of cells that migrate from the ear to the tail while depositing sensory organs in a process requiring dynamic FGF signaling (Chitnis, 2026). FGF/ERK signaling also has roles in adult zebrafish during regeneration: the FGF ligand *fgf20a* is required for regeneration of amputated fins, and scale regeneration is regulated by ERK signaling waves (Bangru et al., 2025). Finally, dysregulation of FGF/ERK signaling has a wide range of pathological consequences, including cancer and mosaic RASopathies, that can be modeled in zebrafish (Garlisi Torales et al., 2024; Monroe et al., 2021).

Strategies to dynamically manipulate FGF/ERK signaling *in vivo* with spatiotemporal precision are therefore valuable for mechanistic dissection of a diverse range of conserved biological processes. Molecular optogenetic approaches (Kolar et al., 2018) can enable precise experimental control over signaling by coupling light exposure to pathway activation (Farahani et al., 2021; Rogers and Müller, 2020). Light exposure is straightforward to control and allows researchers to determine when, where, at what level, and how long to activate signaling. This provides opportunities to experimentally manipulate signaling gradients, levels, and pulses to investigate their functions. The zebrafish embryo is ideal for molecular optogenetic approaches because it is a translucent, externally fertilized, established vertebrate model system with robust molecular, genetic, and genomic resources (Bedell et al., 2025).

Recently, we developed a zebrafish-optimized blue light-responsive optogenetic FGF/ERK activator, which we refer to as “bOpto-FGF” for **<u>b</u>**lue-light **<u>Opto</u>**genetic **<u>FGF</u>** signaling activator (Fig. 1A, Addgene #232639) (Iannucci et al., 2026). Binding of FGF ligand to receptor tyrosine kinases (RTKs) leads to RTK dimerization and phosphorylation, subsequently activating phosphorylation of multiple effectors including ERK (Ornitz and Itoh, 2015). This signaling cascade regulates gene expression and activates feedback regulators. Using the chassis described in (Sako et al., 2016), we fused the zebrafish *fgfr1a* RTK intracellular kinase domain to the blue light-dimerizing light oxygen voltage-sensing (LOV) domain AUREOCHROME from the algae *Vaucheria frigida* (VfAU1) (Takahashi et al., 2007; Toyooka et al., 2011). The fusion protein is tethered to the plasma membrane with a myristoylation motif to mimic the endogenous RTK environment and includes a C-terminal HA tag for immunofluorescence detection. Blue light exposure leads to dimerization of the LOV-containing fusion proteins, initiating RTK phosphorylation and FGF signaling activation (Fig. 1A). We previously introduced bOpto-FGF into zebrafish embryos by injecting mRNA at the one-cell stage and systematically assessed its properties at early gastrulation (Iannucci et al., 2026). Injected embryos robustly activate ectopic ERK phosphorylation and FGF target gene expression in response to blue light ∼455 nm, but are not responsive to wavelengths above 495 nm. The level of signaling activation is dependent on light dosage, and on/off kinetics are rapid: Ectopic ERK phosphorylation is evident within 2 minutes of light exposure and returns to baseline about 10 minutes after light removal. We also demonstrated spatially localized signaling activation in mRNA-injected embryos.

**Fig. 1:**
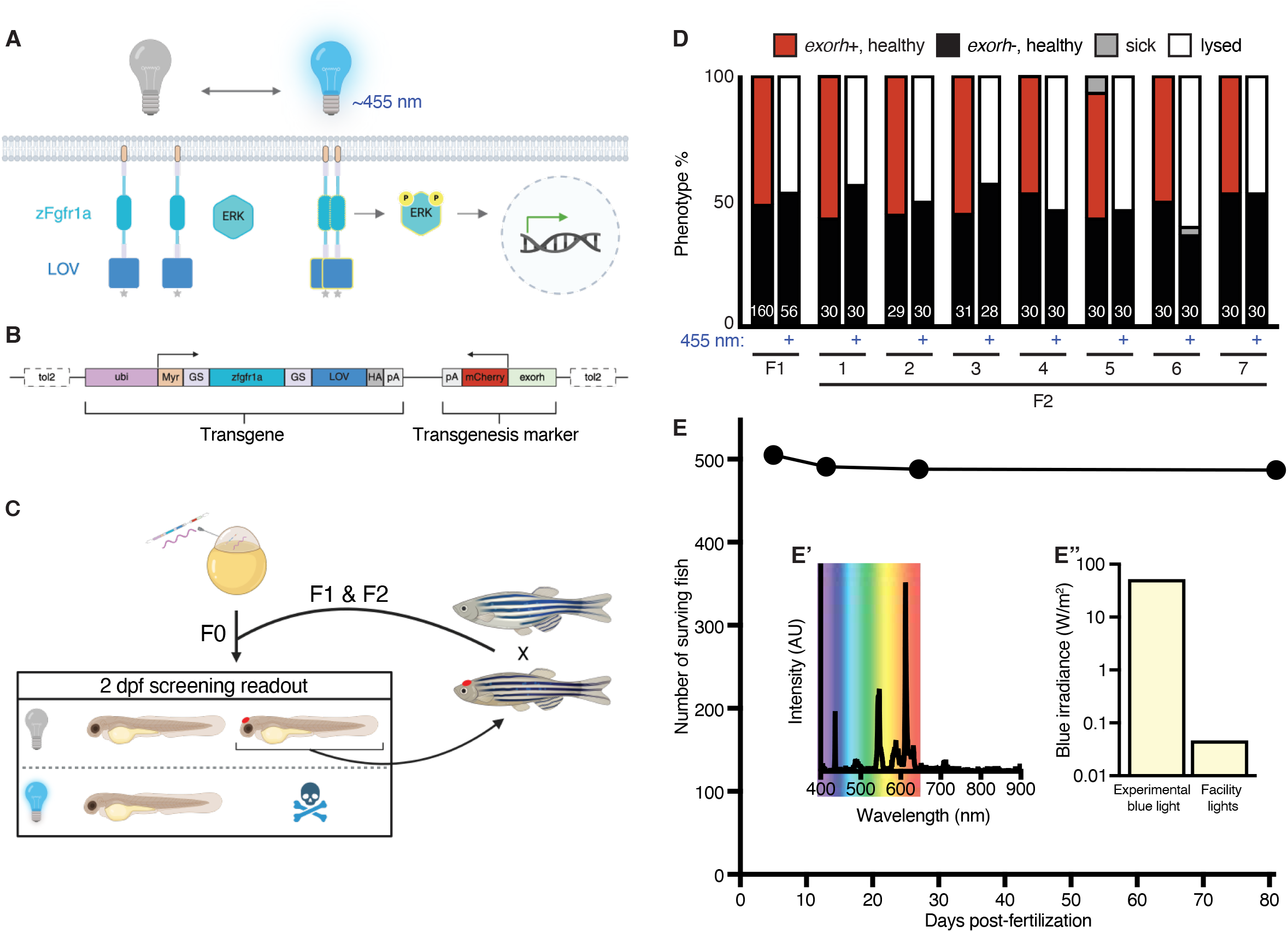
Generation of the *Tg(ubi:bOpto-FGF)* line. A) bOpto-FGF strategy to activate FGF/ERK signaling. Blue light exposure leads to dimerization of LOV-RTK fusion proteins, activating ERK phosphorylation and signaling. B) Schematic of the *ubi:bOpto-FGF* transgene cassette. ubi = ubiquitin promoter, Myr = myristoylation motif, GS = glycine/serine linker, LOV = light oxygen voltage-sensing domain, HA = hemagglutinin tag, pA = polyA tail, exorh = pineal gland promoter. C) Transgenic screening strategy. Injected F0s were exposed to blue light or dark starting at 1.5-2 hours post-fertilization. At 2 days post-fertilization (dpf) embryos were screened for phenotypes consistent with ectopic FGF signaling (lysis) and expression of the mCherry pineal gland marker. This strategy was repeated for F1 and F2 generations. D) Outcross progeny from an F1 founder and seven of its F2 descendants were either reared in the dark or exposed to 455 nm light and scored at 2 dpf. White numbers indicate number of scored embryos. E) Survival of *Tg(ubi:bOpto-FGF)* heterozygotes was tracked starting at 5 dpf and assessed at the indicated time points. Insets show spectrum of facility lights (E’), and irradiance between 420-525 nm from these lights versus 455 nm LEDs used in uniform exposure experiments (E”).

To date, bOpto-FGF and other optogenetic FGF/ERK tools have mainly been introduced into zebrafish embryos by injecting mRNA (Benman et al., 2022; Iannucci et al., 2026; Patel et al., 2025; Patel et al., 2019) (or DNA (Kainrath et al., 2017)). Although useful in many experimental contexts, the mRNA injection strategy poses several challenges. First, successful injections require significant training, time, reagents, and equipment including needles, injection rigs, and synthesized mRNA (Saul et al., 2023). Second, injected mRNA does not always distribute uniformly through the embryo (Iannucci et al., 2026), leading to spatially heterogenous signaling responses that could complicate experiment interpretation. Third, mRNA degrades over time, meaning that light responsiveness will eventually wane, preventing experiments at later developmental stages.

To overcome these challenges, here we introduce and characterize a novel transgenic zebrafish, *Tg(ubi:bOpto-FGF)*, that expresses bOpto-FGF via a ubiquitous promoter. We show that ERK phosphorylation is robustly activated with blue light exposure in heterozygous and homozygous embryos with greater spatial uniformity compared to mRNA-injected embryos, characterize changes in ectopic ppERK levels over time in response to continuous light exposure, observe light-dependent FGF/ERK signaling activation starting at blastula stages through 24 hours post-fertilization (hpf), and demonstrate spatially-localized FGF/ERK signaling activation in the head, trunk, and tail at 1 day post-fertilization (dpf). In contrast to a previously characterized periderm-specific optogenetic line (*Tg(krt4:Dronpa-MEK203)^pd404^*(Ramkumar et al., 2025)), *Tg(ubi:bOpto-FGF)* expresses a zebrafish-optimized tool driven by a ubiquitous promoter, enabling whole-embryo spatiotemporal manipulations. This novel transgenic line is a valuable resource for zebrafish researchers studying dynamic FGF/ERK signaling in diverse biological processes including germ layer patterning, organogenesis, regeneration, and disease.

## RESULTS

### Strategy to generate transgenic zebrafish ubiquitously expressing bOpto-FGF

To maximize potential applications of bOpto-FGF, we created a transgenic zebrafish line with this optogenetic tool driven by a ubiquitous promoter. We reasoned that desired tissues and stages could be targeted by controlling when and where light exposure occurs, given that temporally and spatially restricted signaling activation is possible in *bOpto-FGF* mRNA-injected embryos (Iannucci et al., 2026). Below we describe the derivation of the line to provide guidance in scenarios where shipping restrictions complicate line sharing.

We used the Tol2 transgenesis system (Felker and Mosimann, 2016; Kawakami, 2007; Kwan et al., 2007) to generate F0 zebrafish harboring a transgene containing the *ubi* promoter described in (Mosimann et al., 2011) driving bOpto-FGF expression, as well as a transgenesis marker (Fig. 1B) (Addgene #260611). The 3.5 kb *ubi* promoter region is upstream of the zebrafish *ubiquitin B* locus and is reported to drive ubiquitous expression in nearly all cell types from embryonic stages through adulthood. We cloned this promoter in front of bOpto-FGF followed by a polyadenylation signal. To aid in identifying transgene-positive fish, we included a transgenesis marker in the opposite orientation comprised of the pineal gland-specific *exorh* promoter driving mCherry, followed by a polyadenylation signal (Kemmler et al., 2023). The highly localized pineal gland *exorh* promoter was chosen to minimize overlap with RFP reporter lines that may be used in downstream applications. mCherry was selected because its ∼580 nm excitation wavelength does not activate bOpto-FGF (Iannucci et al., 2026), allowing for the screening of pineal gland RFP-positive embryos without inadvertently activating FGF/ERK signaling. Marker expression is easily detectable at 2 days-post fertilization (dpf). To mediate insertion into the genome, these elements were flanked by Tol2 sites.

Wild-type AB embryos at the one-cell stage were injected with 25 pg of this plasmid and 25 pg of *Tol2* mRNA. At ∼2 hours post-fertilization (hpf), healthy embryos were split into two groups and either kept in the dark or exposed to blue light (as described in (Saul et al., 2023) and (Iannucci et al., 2026); unless otherwise noted, irradiance and wavelength were 50 W/m^2^ and 455 nm, respectively, here and in all subsequent experiments.). At 2 dpf, embryos from both groups were screened for 1) mCherry expression in the pineal gland indicating transgene integration, and 2) phenotypes consistent with ectopic FGF signaling to assess bOpto-FGF activity (Fig. 1C). Strong ectopic FGF/ERK signaling leads to developmental defects or more commonly lysis by 1 dpf (Iannucci et al., 2026). mCherry-positive injected F0s were raised to adulthood and outcrossed to wild-type fish, and the resulting F1 embryos were screened as described above. This process was repeated to generate F2s and identify a stable line.

We identified an F1 fish and its F2 progeny that consistently produced outcross progeny demonstrating two ideal qualities (Fig. 1D). First, in the dark group, ∼50% of embryos were healthy and positive for the transgenesis marker, suggesting negligible “dark leakiness” (i.e., ectopic activation in the dark). Second, in the light-exposed group, ∼50% of embryos lysed, and the remaining embryos were transgene-negative, suggesting robust light-mediated signaling activation. The ∼50% transmission rate of the transgene is consistent with a single insertion. The *Tg(ubi:bOpto-FGF;exorh:mCherry)^y724^*line characterized here was derived from this F1 founder and is hereafter referred to as *Tg(ubi:bOpto-FGF)*.

### Tg(ubi:bOpto-FGF) husbandry

The *ubi* promoter drives nearly uniform expression in embryonic and adult zebrafish (Mosimann et al., 2011), and bOpto-FGF activates ectopic FGF/ERK signaling at ∼455 nm (Iannucci et al., 2026), a wavelength that can be emitted by standard overhead vivarium “white light” sources. It was therefore unclear whether *Tg(ubi:bOpto-FGF)* fish would require special husbandry, for example housing in a light cabinet equipped with > 495 nm light sources. We therefore monitored survival of 505 *Tg(ubi:bOpto-FGF)* heterozygous fry starting at 5 dpf when they were introduced into our fish facility (Fig. 1E). 97.2% of the fish reached adulthood with standard husbandry care and light cycles. We also measured the spectra and irradiance of the ambient light in our facility, which contained wavelengths across the visible light spectrum (∼400-700 nm), including “blue” (420-525 nm) light that could potentially activate bOpto-FGF (Fig. 1E’). However, the measured irradiance within the “blue” range was relatively low, particularly in comparison to the irradiances used in our experimental activation (Fig. 1E”). Under these conditions, *Tg(ubi:bOpto-FGF)* fish reliably breed.

### Robust light-mediated FGF target gene expression and spatially uniform FGF/ERK signaling activation in *Tg(ubi:bOpto-FGF)* embryos

Although *Tg(ubi:bOpto-FGF)* embryos exposed to continuous blue light starting at 2 hpf typically lyse by 2 dpf (Fig. 1D) consistent with ectopic FGF/ERK activation, lysis is a non-specific phenotype. We therefore sought to directly assess FGF/ERK activation by quantifying ppERK levels. To determine whether *Tg(ubi:bOpto-FGF)* heterozygous and homozygous embryos activate FGF/ERK signaling in response to blue light exposure, and to compare the activity of this line with mRNA-injected embryos, we exposed transgenic embryos and embryos injected with *bOpto-FGF* + *GFP* mRNA to 455 nm light for 30 min starting at early gastrulation (shield stage, ∼7 hpf). GFP was included in injection mixes to monitor distribution of mRNA (Iannucci et al., 2026). Homozygous embryos were obtained from incrosses of homozygous parents, while heterozygotes were obtained from outcrosses of homozygous males to wild-type females. Exposed embryos and dark controls were fixed, and hybridization chain reaction immunofluorescence (HCR-IF) was used to detect ppERK (Iannucci et al., 2026). ppERK-intensity fold changes were then calculated compared to respective dark controls (see Methods). For this experiment and all that follow, three independent biological replicates were performed with 7-8 embryos analyzed per condition (except where noted).

Transgenic and mRNA-injected embryos all showed a strong, significant increase in ppERK levels in response to blue light exposure compared to respective dark controls (*p* <0.0001, Fig. 2A,C; Fig. S1, S2; Tables S1-S2), demonstrating that *Tg(ubi:bOpto-FGF)* enables light-mediated activation of FGF/ERK signaling. ppERK fold changes were not significantly different between light-exposed mRNA-injected embryos and *Tg(ubi:bOpto-FGF)* homozygotes (*p* = 0.9986); however, the fold change in light-exposed *Tg(ubi:bOpto-FGF)* heterozygotes was significantly lower than that in exposed mRNA-injected embryos *and* homozygotes (both *p* < 0.0001).

**Fig. 2:**
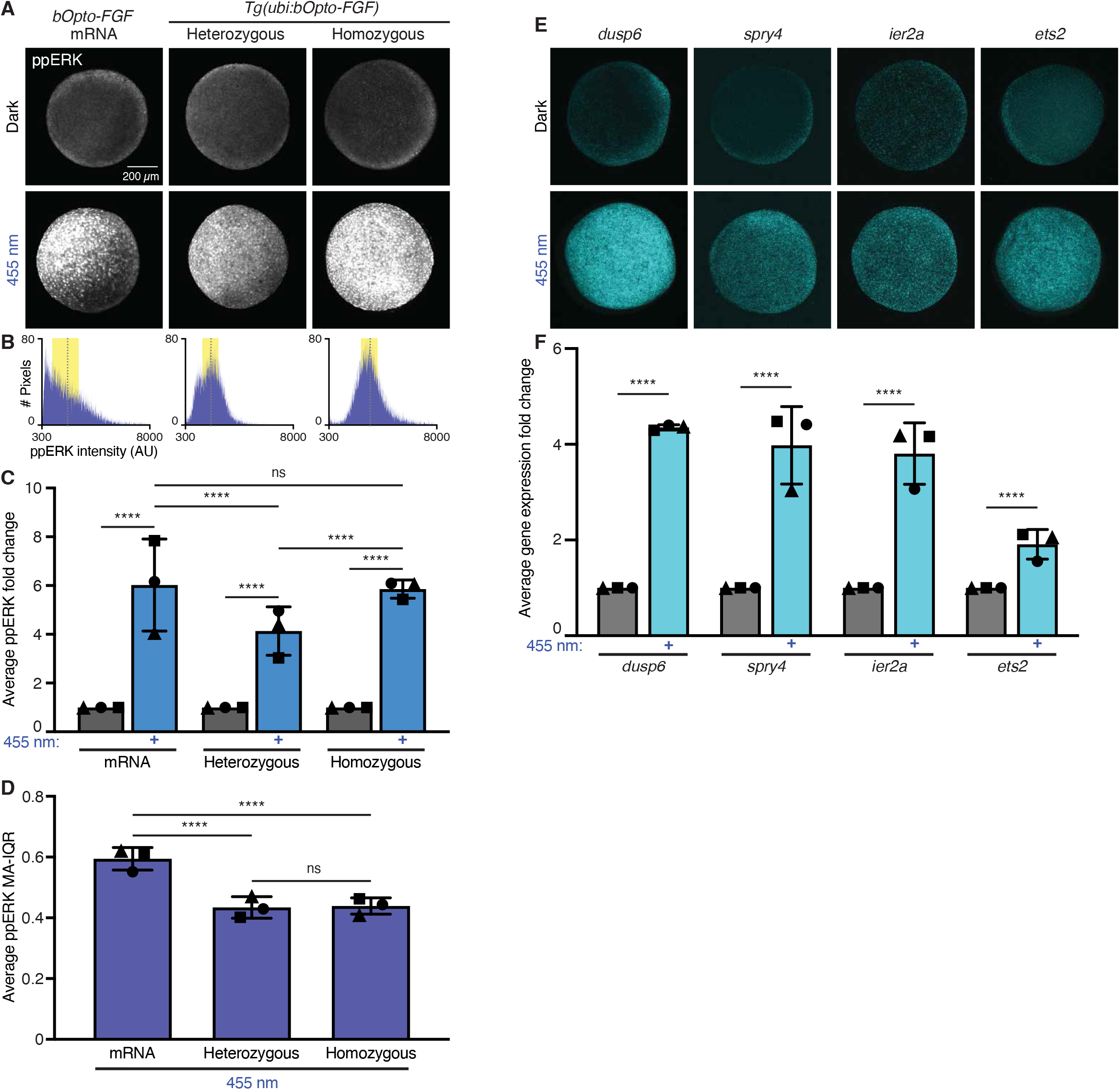
Optogenetic FGF/ERK signaling & target-gene activation. (A-D) *Tg(ubi:bOpto-FGF)* and bOpto-FGF + GFP mRNA-injected embryos were exposed to 455 nm light for 30 min beginning at shield stage. Embryos were fixed alongside unexposed controls. HCR-IF staining for ppERK was used to quantify FGF/ERK signaling. A) ppERK staining in representative embryos. B) Histograms of ppERK pixel intensities from exposed images in (A). Gray dotted lines indicate thresholded means, yellow shading indicates interquartile range (IQR). See Fig. S1 for more examples. C) Intensity fold changes. Each symbol represents averaged data from one repliacte. See Fig. S2 for individual replicates. D) Average ppERK mean-adjusted IQR (MA-IQR) from the three replicates above. (E-F) *Tg(ubi:bOpto-FGF)* heterozygous embryos were exposed to 455 nm light for 45 min beginning at 50% epiboly. Embryos were fixed alongside unexposed controls. HCR-FISH staining for the indicated genes was performed. E) Representative embryos shown. F) HCR-FISH intensity fold changes. Each symbol represents averaged data from one replicate. See Fig. S3 for individual replicates. Error bars show SD. Scale bar = 200 µm.

We have previously noted non-uniform ectopic FGF/ERK activation in some light-exposed *bOpto-FGF* mRNA-injected embryos due to inhomogeneous mRNA distribution (Fig. S1) (Iannucci et al., 2026). To compare the spatial uniformity of FGF/ERK activation in transgenic and mRNA-injected embryos, we calculated the mean-adjusted interquartile range (MA-IQR) of ppERK pixel intensities for each individual light-exposed embryo (Fig. 2B,D; Fig. S1, S2; Tables S1 and S3). Briefly, pixel intensities were arranged from lowest to highest and visualized as histograms. The difference between the 75^th^ and 25^th^ percentiles of this data was calculated as the IQR, a common measure of spread. To normalize for differences in absolute intensity between embryos, each embryo’s IQR was then divided by its mean ppERK pixel intensity. Lower MA-IQR values indicate a narrower distribution of ppERK intensities, consistent with more spatially uniform signaling activation. *Tg(ubi:bOpto-FGF)* homozygotes and heterozygotes both had significantly lower average MA-IQR values compared to mRNA-injected embryos (both *p* < 0.0001). Average MA-IQR values did not differ significantly between homozygous and heterozygous embryos (*p* = 0.9852). Based on this comparison, we conclude that light-mediated FGF/ERK signaling activation in *Tg(ubi:bOpto-FGF)* embryos is more spatially uniform compared to mRNA-injected embryos.

As an additional readout for FGF/ERK signaling activation, we used hybridization chain reaction fluorescence *in situ* hybridization (HCR-FISH) (Choi et al., 2018) to assess expression of known FGF target genes *spry4* (Fürthauer et al., 2001), *dusp6* (Tsang et al., 2004), and *ier2a* (Hong and Dawid, 2009), as well as *ets2*, a member of the zebrafish ETS family (Liu and Patient, 2008) that we hypothesized might be positively regulated by FGF signaling similar to other ETS family members (Raible and Brand, 2001; Roehl and Nusslein-Volhard, 2001). Heterozygous *Tg(ubi:bOpto-FGF)* embryos were either kept in the dark or exposed to 455 nm light for 45 min starting at early gastrulation (50% epiboly, ∼6.25 hpf) and subsequently fixed. We chose a 45 min exposure to allow time for transcript accumulation. HCR-FISH showed significant upregulation of all four genes in light-exposed embryos compared to dark controls (*p* < 0.0001, Table S4) (Fig. 2E-F; Fig. S3).

Together, these analyses demonstrate robust, spatially uniform blue light-mediated activation of FGF/ERK signaling and target gene expression in *Tg(ubi:bOpto-FGF)* embryos during early gastrulation.

### Rapid, long-lasting light-mediated FGF/ERK signaling activation in *Tg(ubi:bOpto-FGF)*

FGF signaling can activate negative feedback (Ornitz and Itoh, 2015). To determine whether ectopic signaling levels change over time during light exposure, we exposed late-blastula stage (30% epiboly) heterozygous *Tg(ubi:bOpto-FGF)* embryos to continuous 455 nm light, fixed embryos at 3, 7, 15, 39, 63, and 79 minutes into exposure, and quantified ppERK using HCR-IF (Fig. 3; Fig. S4; Tables S1, S5A, and S5B). We chose to perform these and subsequent experiments in heterozygotes (obtained from homozygous males crossed to wild-type females), anticipating that users of *Tg(ubi:bOpto-FGF)* will outcross it to mutant or reporter lines and may therefore prefer to perform experiments in heterozygotes. Time points were selected based on assessment of prior *bOpto-FGF* mRNA-injection experiments (Iannucci et al., 2026). We then calculated fold changes in ppERK levels compared to time-matched dark controls. (Dark controls collected at 10 minutes served as controls for both the 3- and 7-minute exposed time points because we expect no meaningful difference in endogenous FGF/ERK signaling over this short time frame. 3, 7, 10, and 39-minute timepoints were imaged on one day with the same imaging settings. 15, 63, and 79-minute timepoints were imaged the next day with different settings. These differences are accounted for by our fold change calculation strategy.)

**Fig. 3:**
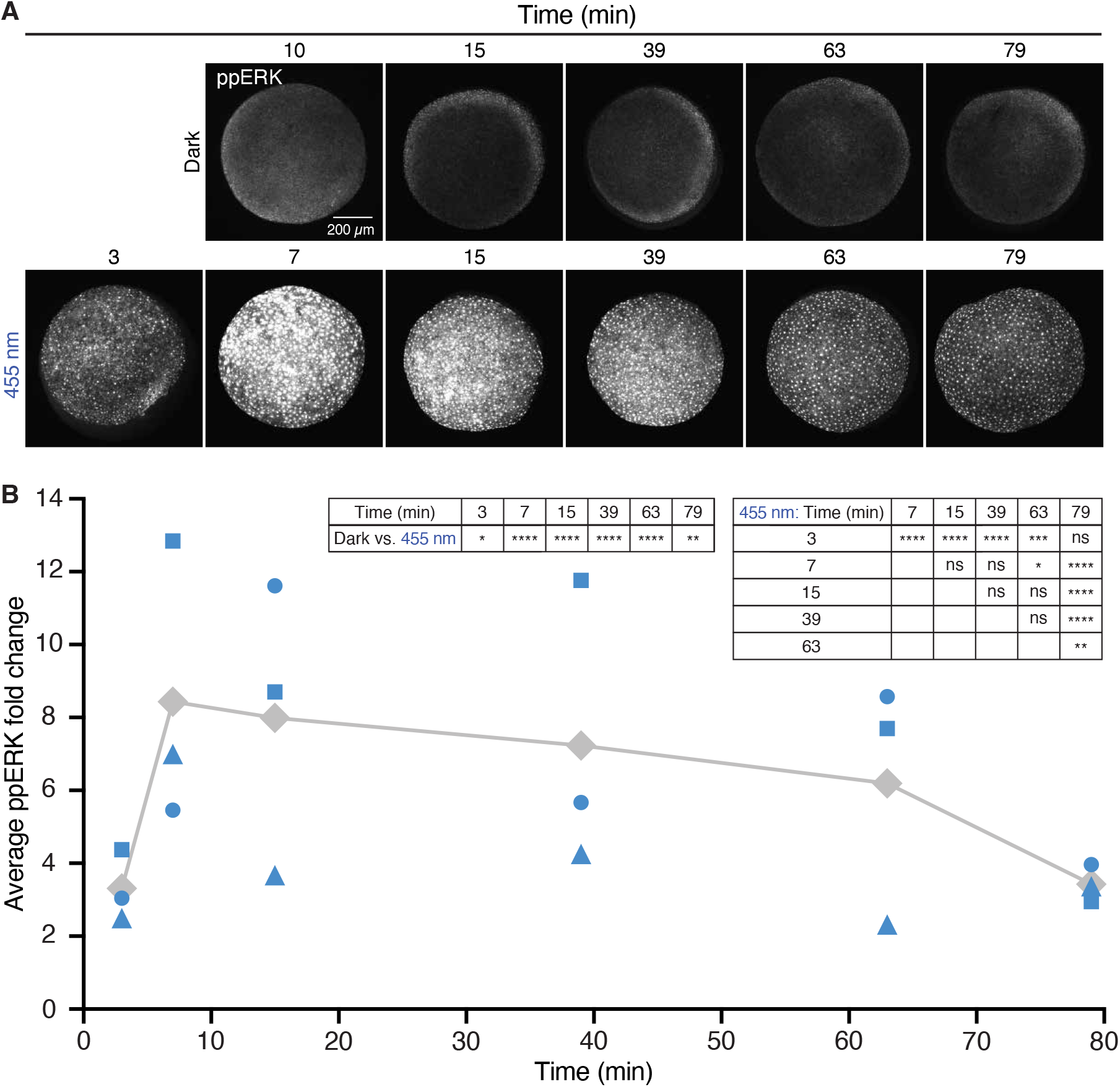
*Tg(ubi:bOpto-FGF)* signaling activation kinetics. Heterozygous *Tg(ubi:bOpto-FGF)* embryos were exposed to continuous 455 nm light beginning at 30% epiboly. Embryos were removed and fixed along with time-matched unexposed controls at the indicated durations. A) HCR-IF ppERK staining in representative light-exposed embryos and time-matched unexposed controls. 3, 7, 10, and 39-minute timepoints were imaged on one day with the same imaging settings. 15, 63, and 79-minute timepoints were imaged the next day with different settings. Scale bar = 200 µm. B) ppERK intensity fold changes. Each blue symbol represents averaged data from one independent replicate. Gray diamonds represent time point means. See Fig. S4 for data from individual replicates.

At all time points tested, light-exposed embryos showed significantly higher levels of ppERK compared to dark controls (Table S5A). At 3 minutes of light exposure, signaling activation appeared highly variable compared to other time points, and the fold change in ppERK was significantly lower than the fold changes at 7, 15, 39, and 63 minutes (Fig. S4; Table S5B). The ppERK fold change at 79 minutes was significantly lower than fold changes at 7, 15, 39, and 63 minutes. These results show that continuous light exposure starting at 30% epiboly can activate FGF/ERK signaling within 3 minutes, and that ectopic signaling is still activated at 79 minutes but at lower levels compared to earlier time points, suggesting possible negative feedback.

### *Tg(ubi:bOpto-FGF)* heterozygotes are light-responsive starting at blastula stages through at least 24 hours post-fertilization

Developmental processes regulated by dynamic FGF/ERK signaling begin at blastula stages and continue beyond 24 hpf (Chitnis, 2026; Gibbs et al., 2017; Jones and Mullins, 2022; Riley, 2021). To determine whether *Tg(ubi:bOpto-FGF)* can reliably activate FGF/ERK signaling during these developmental stages, we exposed embryos to 455 nm light for 30 minutes starting at five stages: sphere, 30% epiboly, shield, bud, and 24 hpf (Kimmel et al., 1995). Exposed embryos and stage-matched dark controls were fixed immediately after the 30 minutes and HCR-IF was performed to quantify changes in ppERK levels (Fig. 4; Fig. S5; Tables S1, S6A, and S6B). Experiments were performed in heterozygous embryos that were progeny of male homozygotes outcrossed to wild-type females.

**Fig. 4:**
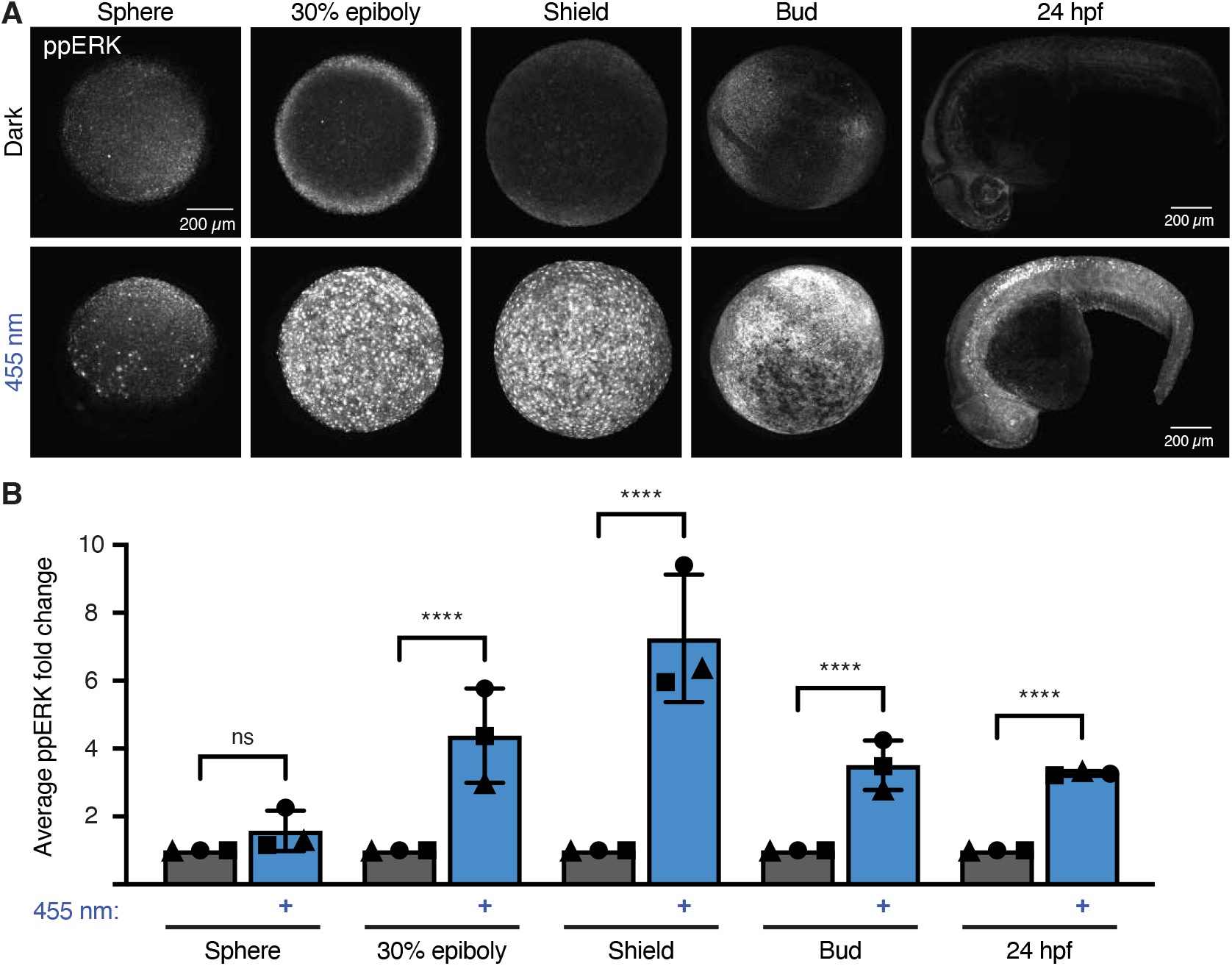
Light-mediated activation of FGF/ERK signaling from blastula stages thorough 24 hours post-fertilization (hpf). Heterozygous *Tg(ubi:bOpto-FGF)* embryos were exposed to 455 nm light beginning at the indicated stages. Embryos were fixed after 30 min along with stage-matched unexposed controls. A) HCR-IF staining for ppERK in representative embryos. Scale bar = 200 µm. B) ppERK intensity fold changes. Each symbol represents the average from one replicate. Error bars show SD. See Fig. S5 for data from individual replicates.

At sphere stage, the earliest time point tested, ppERK intensity was not significantly different between exposed and unexposed embryos overall (*p* = 0.5748, Table S6A). However, some exposed embryos appeared to have ectopic ppERK, suggesting that heterozygous embryos may begin to become light sensitive around or after sphere stage (Fig. S5). Activation at sphere stage is therefore highly variable and not reliable in male-derived heterozygotes. In contrast, at the next tested time point, 30% epiboly, ppERK levels were significantly and strongly increased in exposed embryos compared to dark controls (*p* < 0.0001). ppERK levels were also significantly increased in exposed embryos compared to dark controls at shield, bud, and 24 hpf (*p* < 0.0001). At 24 hpf, we detected ectopic FGF/ERK signaling across light-exposed embryos. To assess *bOpto-FGF* mRNA distribution at 24 hpf, we performed HCR-FISH targeting *LOV*, the light-sensitive domain in bOpto-FGF which is not present in the wild-type zebrafish genome. Consistent with the ectopic FGF/ERK signaling observed across light-exposed transgenic embryos at 24 hpf, we detected the embryo-wide presence of *LOV* mRNA, although it appears to be enriched in somites (Fig. S6) (Note: LOV HCR-FISH replicate 1 had 4 embryos per condition in contrast to the 8 used in replicates 2 and 3). Together, our results show that *Tg(ubi:bOpto-FGF)* heterozygotes derived from male outcrosses become light responsive by 30% epiboly and continue through at least 24 hpf.

### Spatially localized FGF/ERK signaling activation in *Tg(ubi:bOpto-FGF)*

One advantage of optogenetic strategies is the ability to spatially control activation by restricting light exposure to a desired area. Localized ectopic FGF/ERK activation has been demonstrated in *bOpto-FGF* mRNA-injected embryos at gastrulation stages (Iannucci et al., 2026). To determine whether *Tg(ubi:bOpto-FGF)* enables spatially localized activation at later stages, we exposed the head, trunk, or tail of 1 dpf heterozygotes to a 170 x 170 µm square of 455 nm LED light (11.42 W/m^2^) for 1 minute using a digital micromirror device. HCR-IF staining demonstrated elevated levels of ppERK at the targeted regions (Fig. 5; Fig. S6). *Tg(ubi:bOpto-FGF)* can therefore be used for applications requiring spatially restricted ectopic FGF/ERK activation.

**Fig. 5.**
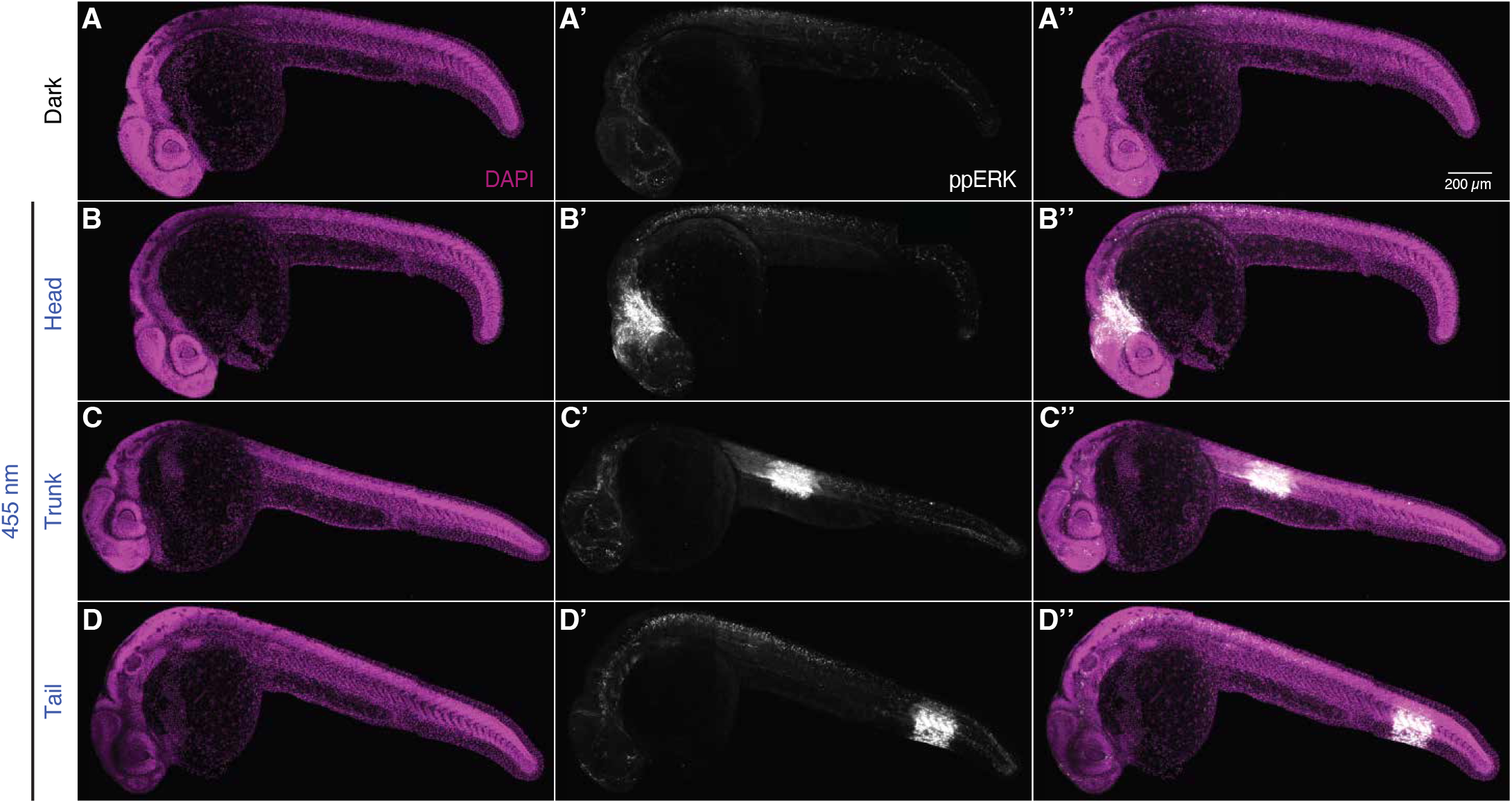
Spatially localized activation of FGF/ERK signaling. Heterozygous *Tg(ubi:bOpto-FGF)* embryos were raised in the dark until 1 dpf, then exposed to a 170 x 170 µm square of 455 nm light for 1 min using a digital micromirror device mounted on a confocal microscope. Representative DAPI (magenta, A, B, C, D) and ppERK (gray, A’, B’, C’, D’) images as well as merged images (A”, B”, C”, D”) are shown for dark controls (A-A”) and head-exposed (B-B”), trunk-exposed (C-C”), and tail-exposed (D-D”) embryos. See Fig. S6 for additional images from three biological replicates. Scale bar = 200 µm. Images from replicate 2.

## DISCUSSION

### A novel transgenic for robust optogenetic activation of FGF/ERK signaling in zebrafish

The ability to dynamically manipulate FGF/ERK signaling *in vivo* creates unprecedented experimental opportunities to study germ layer patterning, organogenesis, regeneration, disease, and more. Here, we introduce a novel transgenic zebrafish, *Tg(ubi:bOpto-FGF;exorh:mCherry)^y724^*, that enables rapid optogenetic activation of FGF/ERK signaling starting at late-blastula stages through at least 24 hpf (Fig. 4, Fig. S5), and provide guidance (Fig. 1) and reagents to re-derive the line in cases where shipping is challenging (Addgene #260611, #260612). We chose to drive bOpto-FGF expression with a ubiquitous promoter to allow users to flexibly manipulate signaling in the desired tissue and stage by spatiotemporally controlling light exposure, and demonstrated rapid (within 1 minute), specific ectopic activation in the head, trunk, or tail at 1 dpf using this strategy (Fig. 5; Fig. S6). In addition, we performed many of our characterizations in heterozygotes, reasoning that users may be interested in outcrossing to mutant or reporter lines, such as the recently optimized “modERK-KTR” zebrafish FGF/ERK signaling reporter (Wilcockson et al., 2023). The *exorh:mCherry* transgenesis marker was also chosen for this reason, because its expression is restricted to a small gland unlikely to conflict with other RFP reporters (Kemmler et al., 2023). We note that homozygous adults are viable, and that shield-stage homozygotes show stronger signaling activation compared to heterozygotes (Fig. 2; Fig. S1, S2). We speculate that this is due to expression from two copies rather than one copy of the transgene, which, based on the ∼50% segregation upon outcrossing (Fig. 1D), appears to be a single insertion. Below we discuss some advantages, limitations, recommendations, and future directions for this transgenic strategy.

### Advantages of *Tg(ubi:bOpto-FGF)* compared to mRNA injection

bOpto-FGF has previously been introduced into zebrafish by injecting mRNA at the one-cell stage (Iannucci et al., 2026). Although mRNA injection is ideal for many experimental designs, our novel transgenic can reduce experimental friction by obviating the time, training, reagents, and equipment necessary to perform injections (Saul et al., 2023). At early gastrulation, levels of ectopic ppERK did not differ significantly between light-exposed mRNA-injected embryos and homozygous transgenics, but were lower in exposed heterozygous transgenics compared to either (Fig. 2C; Fig. S2). Also at this stage, light-activated FGF/ERK signaling was more spatially uniform in transgenic embryos compared to mRNA-injected embryos (Fig. 2B,D; Fig. S1, S2). This decreased variability may be valuable in experiments in which identical responses from all exposed cells are desired. We also determined similarly rapid signaling activation in transgenics compared to mRNA-injected embryos: Transgenic heterozygotes showed activation of signaling within 3 minutes of exposure at 30% epiboly (Fig. 3; Fig. S4), the earliest time point tested in this assay, and mRNA-injected embryos at shield stage showed activation within 2 minutes of exposure (Iannucci et al., 2026). In addition, our spatially localized activation experiments at 1 dpf showed ectopic FGF/ERK activation in the targeted region after only 1 minute of light exposure (Fig. 5, Fig. S6). Finally, the *ubi* promoter is reported to drive expression in most cells from embryonic stages through adulthood (Mosimann et al., 2011). In contrast, transiently injected mRNA will eventually degrade. Here we have shown that *Tg(ubi:bOpto-FGF)* heterozygotes enable light-mediated FGF/ERK signaling activation starting at 30% epiboly until at least 24 hpf (Fig. 4, Fig. S5).

### *Tg(ubi:bOpto-FGF)* husbandry and handling

In contrast to mRNA-injected embryos, the *Tg(ubi:bOpto-FGF)* line presents a potential husbandry challenge: inadvertent blue light exposure leading to deleterious FGF/ERK activation. However, *Tg(ubi:bOpto-FGF)* fish survive and breed in our facility, even though our facility’s white light source contains emissions in the 420-525 nm range (Fig. 1E’). Importantly, we caveat that different lighting conditions could lead to problematic ectopic activation. In facilities where lighting conditions are determined to be detrimental, fish could be housed in a light cabinet emitting wavelengths above 495 nm which do not activate bOpto-FGF (Iannucci et al., 2026). Alternatively, a novel bipartite transgenic strategy could be designed using GAL4/UAS, Cre/Lox, or QF/QUAS to avoid potential challenges arising from light sensitive adults.

Our work suggests that *Tg(ubi:bOpto-FGF)* embryos can be handled in standard room lighting conditions without obvious deleterious signaling activation for the first several hours of development. Heterozygous sphere stage *Tg(ubi:bOpto-FGF)* overall did not significantly activate ectopic FGF/ERK signaling when exposed to 455 nm light for 30 minutes; however, we did observe some light-exposed embryos at this stage that appeared to have elevated ppERK levels (Fig. 4; Fig. S5). We therefore recommend treating embryos older than high stage as light sensitive. Older embryos can be protected from inadvertent FGF/ERK signaling by wrapping dishes in foil, or by handling embryos in a room with red light sources (above 495 nm) and using red filter paper to cover dissecting microscope light sources, similar to the handling recommendations for mRNA-injected embryos (Iannucci et al., 2026).

### Future directions

Future work could further expand the utility of *Tg(ubi:bOpto-FGF)*. Although we expect the light dosage sensitivity, off kinetics, pathway specificity, and wavelength dependence that were determined previously for bOpto-FGF in mRNA-injected embryos are similar (Iannucci et al., 2026), characterization of these properties in the transgenic might be useful. In addition, our evidence suggests that *Tg(ubi:bOpto-FGF)^y724^* is a single transgene insertion in an unknown locus (Fig. 1D). If needed, the insertion site could be identified using Tol2 insertion mapping strategies (Meng et al., 2025; Wei et al., 2025). Further, we found that heterozygous embryos derived from male outcrosses are not significantly light responsive until after sphere stage (Fig. 4; Fig. S5). Although this is beneficial for handling, it could pose a challenge for experimentalists wishing to manipulate FGF/ERK at very early stages. More detailed characterization of homozygotes, or heterozygotes from female outcrosses, might reveal relevant differences in light responsiveness at earlier stages given that the *ubi* promoter is reported to drive maternal expression (Mosimann et al., 2011). Finally, although we demonstrated spatially localized signaling activation (Fig. 5; Fig. S6), tissue-specific bOpto-FGF lines could further tailor this tool toward highly specific biological questions—as evidenced by the insights into skin development facilitated by the periderm-specific ERK activating line (Ramkumar et al., 2025).

Overall, the *Tg(ubi:bOpto-FGF)* zebrafish is a powerful reagent that can be used to study a wide range of outstanding biological questions, including how different levels and durations of signaling influence cell fate decisions in multiple contexts. Precise spatiotemporal manipulation of FGF/ERK signaling dynamics *in vivo* promises to provide new insights in developmental biology, regenerative biology, and translational research.

## Supporting information

Anderson et al supplemental material

## Acknowledgements

We thank Brant Weinstein and Harry Burgess and their labs for advice and reagents, and members of the Rogers lab and Jeffrey Farrell and his lab for helpful discussions, feedback, and reagents. We especially thank Velanganni Selvaraj Maria Thomas for help with experiments and statistics, and Micaela Murphy for designing the *ier2a* and *ets2* HCR-FISH probes. We are grateful to the NIH zebrafish facility staff for maintaining our fish stocks and for discussing facility lighting. Some figures were created in BioRender (Rogers, K. (2026) https://BioRender.com/8t4aenj).

## MATERIALS & METHODS

### Zebrafish husbandry

Zebrafish husbandry and research protocols were approved by the NICHD Animal Care and Use Committee in accordance with the Guide for the Care and Use of Laboratory Animals of the National Institutes of Health. Transgenics were generated in the AB zebrafish (*Danio rerio*) wild-type background. Embryos were incubated at 28-28.5 °C in embryo medium kept at a pH of 7.0-7.4 (reverse osmosis water, “Instant Ocean Sea Salt” (0.25 g/L, Cat. No. SS1-160P), and NaHCO_3_ (∼0.06 g/L, depending on initial pH)).

### *bOpto-FGF* mRNA injections

bOpto-FGF pCS2+ plasmid (Addgene #232639) was linearized with NotI-HF (New England Biolabs Cat. No. R3189L) and transcribed using an SP6 mMessage mMachine kit (Invitrogen AM1340). 1 nl of injection mix containing 3.5 pg *bOpto-FGF* mRNA + 20 pg *GFP* mRNA, and phenol red tracer was injected through the chorion at the one-cell stage ((Iannucci et al., 2026), GFP modified from (Lim et al., 2009)). Embryos were incubated at 28-28.5 °C immediately after injection. Between 1.5 - 2 hpf, healthy, fertilized embryos were transferred to 6-well dishes (Falcon, Cat. No. 353046) and wrapped in aluminum foil until shield stage. In addition, a subset of embryos was exposed to blue light starting ∼2 hpf and phenotyped, along with dark controls, at 1 dpf to ensure mRNA function as in (Iannucci et al., 2026).

### Safe fish handling

To avoid inadvertent FGF/ERK activation, bOpto-FGF-expressing embryos older than 2 hpf were handled in a windowless room illuminated by red light (A19 9W Equivalent 60W, E26 Red LED Colored Light Bulb, UNILAMP, Cat. 653 No. B0C7YZ4KSY) and visualized using a dissecting scope with the light source covered by red gel filter paper (#E106, Rosco, Cat. No. 110084014805-E106). When not being handled, embryos were maintained in petri dishes or 6-well dishes wrapped in aluminum foil.

Fry and adults were housed under standard conditions in the NIH Zebrafish Facility with a 14 hour on / 10 hour off day-night cycle (Main housing lighting: Sylvania FP54/830/HO/ECO 3000K 54W light bulbs; Auxiliary light source: Phillips F32T8/TL935 Deluxe Neutral White). Spectra of the fish facility lights were measured using a compact spectrometer (Thor Labs, Cat. No. CCS200) with a cosine corrector (Thor Labs, Cat. No. CCSB10), and irradiance was measured using a digital optical power meter (Thor Labs, Cat. No. PM100D) attached to a microscope slide sensor head (Thor Labs, Cat. No. S170C).

### Generation of the *ubi:bOpto-FGF* Tol2 expression vector

To generate a Gateway cloning-compatible middle entry vector containing *bOpto-FGF*, Gibson cloning (NEBuilder, New England Biolabs, Cat No. E2621L) was used to combine a *bOpto-FGF* fragment (Addgene #232639, (Iannucci et al., 2026)) and a pME backbone using the following primers:

Gibson_of_pME_bOpto_fgf_pME_backbone_F:

ccagactacgcataaCACCCAGCTTTCTTGTAC

Gibson_of_pME_bOpto_fgf_pME_backbone_R:

ctccccatggtggcAGCCTGCTTTTTTGTACAAAGTTGGC

Gibson_of_pME_bOpto_fgf_fgf_fragment_F:

gtacaaaaaagcaggctGCCACCATGGGGAGTAGC

Gibson_of_pME_bOpto_fgf_fgf_fragment_R:

gtacaagaaagctgggtgTTATGCGTAGTCTGGTACGTCG

Results were confirmed via sequencing. *pME-bOpto-FGF* (Addgene #260612) was then combined in an LR clonase Gateway reaction as described in the Invitrogen Multisite Gateway Manual (Invitrogen Cat. No. 12538120) with p5E-ubi (Addgene #27320, (Mosimann et al., 2011)), p3E-SV40-polyA (Kwan et al., 2007), and *pDEST-exorh:mCherry* (Addgene # 195989, (Kemmler et al., 2023)) to generate *pExpress-ubi:bOpto-FGF* (Addgene #260611). Results were confirmed via sequencing. See Supplementary Materials for plasmid maps.

### Generating and screening *Tg(ubi:bOpto-FGF;exorh:mCherry)* transgenics

To generate *Tg(ubi:bOpto-FGF;exorh:mCherry)^y724^* transgenics, AB wild-type embryos were injected at the one-cell stage with 1 nl injection mix containing 25 pg *Tol2* transposase mRNA, 25 pg *pExpress-ubi:bOpto-FGF* plasmid, and phenol red tracer. mRNA was synthesized from plasmids linearized with the NotI enzyme (New England Biolabs Cat. No. R3189L) via the SP6 mMessage mMachine kit (Invitrogen AM1340).

For these injected F0 embryos and subsequent generations, the following screening strategy was used with generation-specific modifications to identify promising candidates (Fig. 1C). Between 1.5 and 2 hpf, 30-60 healthy embryos were transferred to a Petri dish and exposed to 455 nm light in an “LED incubator” at 28 °C (Iannucci et al., 2026; Saul et al., 2023). A separate set of 30-60 healthy embryos from the same clutch were transferred to a Petri dish, wrapped in aluminum foil to block light exposure, and placed in the same incubator. At 2 dpf, both groups of embryos were assessed for phenotypes consistent with ectopic FGF signaling (developmental defects and lysis). Embryos were also screened for the presence of the pineal gland transgenesis marker *exorh:mCherry* (Kemmler et al., 2023). Ideal clutches had the following characteristics: 1) a portion of unexposed embryos expressed the transgenesis marker and were healthy (suggesting negligible “dark leakiness”), 2) a similar proportion of embryos exposed to blue light had phenotypes consistent with ectopic FGF signaling, and 3) very few light-exposed, transgenesis marker-positive embryos remained healthy, suggesting that most embryos with the transgene activated ectopic FGF/ERK signaling in response to light.

For the F0 and F1 generations, screening was modified as follows: Due to the random integration mediated by Tol2, embryos in these generations are expected to contain the transgene integrated into different sites. Therefore, selection was more lenient for transgene-positive embryos exhibiting dark leakiness or a lack of response to light. Stringency for these characteristics was higher with F1s than F0s.

For the F2 generation, screening was modified as follows: Maximum stringency was used, and clutches in which ∼50% of unexposed embryos expressed the transgenesis marker (indicating the founder had a single insertion of the transgene) were prioritized. One F1 consistently produced F2 clutches that possessed all desired qualities (Fig. 1D) and was selected to found the *Tg(ubi:bOpto-FGF;exorh:mCherry)^y724^*line characterized here.

To identify heterozygous and homozygous *Tg(ubi:bOpto-FGF)* fish, candidates were outcrossed to wild-types and pineal-gland marker transmission rates were measured. Heterozygotes are expected to have 50% mCherry-positive progeny, whereas homozygotes are expected to have 100%.

### Uniform 455 nm light exposure

Embryos were exposed to uniform 455 nm blue light using the “LED incubators” described in (Saul et al., 2023) and (Iannucci et al., 2026). Pre-sorted embryos in foil-wrapped Petri dishes or 6-well dishes were unwrapped and placed in an LED incubator, with controls still wrapped in foil placed on the shelf below. 50 W/m^2^ irradiance was confirmed prior to all experiments using a digital optical power meter (Thor Labs, Cat. No. PM100D) attached to a microscope slide sensor head (Thor Labs, Cat. No. S170C). At the end of their exposure period, embryos were immediately fixed while still in their respective lighting conditions in 4% formaldehyde and incubated at 4 °C overnight.

### Spatially localized 455 nm light exposure

For spatial activation experiments shown in Fig. 5 and Fig. S6, heterozygous *Tg(ubi:bOpto-FGF)* embryos were reared in the dark until 1 dpf, then manually dechorionated using forceps in a red light-illuminated room (as described in the “Safe fish handling” section above). Embryos were anesthetized using 2 ml of 0.4% MESAB (Ethyl 3-aminobenzoate methanesulfonate, Millipore Sigma cat. No. E10521-10G) in ∼40 ml of embryo medium and transferred to 1.2% low melting agarose (LMA, Invitrogen, Cat. No. 16520100) in embryo medium heated to 42 °C using a glass pipette pump, and mounted on a plastic Petri dish, with one embryo per dish.

To restrict light exposure to a small square, we used conditions similar to those described in (Iannucci et al., 2026). Briefly, an upright laser scanning confocal microscope (Zeiss LSM 800) with a digital micromirror device (DMD, Mightex, Polygon 1000) was used as the illumination source. The light path consisted of the emission from a 455 nm LED (Mightex, Part No. BLS-LCS-0470-14-22) passing through beam combiners in series (Mightex Part No. LCS-BC25-0435, LCS-BC25-0515) to the DMD. Reflected light was directed to the confocal through an infinity port expander (Mightex, Part No. IPX-CHASSIS, IPX-BS-90R10T-UF2) and passed through a W N-Achroplan 10x/0.3 objective before reaching the sample plane. A square ROI was defined using PolyScan4 software (Mightex), and a 640 nm laser (visualized in Zeiss Zen software) was used to position the embryo’s target tissue (head, trunk, or tail) within the ROI. One embryo at a time was exposed to a 170 × 170 μm square of 455 nm LED light at 11.42 W/m^2^ for a duration of 1 min with the output set to 1%. After exposure, embryos were immediately fixed by lifting the agarose pad off the petri dish with a plastic scoop and into a 2 ml Eppendorf tube containing 1 ml 4% formaldehyde. Mounted time-matched unexposed transgenic embryos were also fixed as controls. Embryo ages varied between 24-32 hours post fertilization.

### Hybridization Chain Reaction Immunofluorescence (HCR-IF)

The HCR-IF protocol used here is identical to the protocol described in (Iannucci et al., 2026) based on (Schwarzkopf et al., 2021). As in the referenced protocol, 1:5000 mouse anti-ppERK1/2 primary antibody (Sigma Aldrich #M8159), 1:500 donkey anti-mouse B5 secondary antibody (Molecular Instruments), and B5-647 amplifier (Molecular Instruments) were used to detect ppERK.

### Hybridization Chain Reaction Fluorescence *In Situ* Hybridization (HCR-FISH)

The HCR-FISH protocol used here is identical to the protocol described in (Iannucci et al., 2026) based on (Choi et al., 2018). Probes targeting the *LOV* domain, *ets2*, and the known FGF target genes *dusp6*, *spry4*, and *ier2a* were designed using the probe generator developed in (Kuehn et al., 2022) (see supplementary excel file for sequences). For FGF target gene HCR-FISH experiments, one set of embryos was co-stained for *dusp6* (B2-546 amplifier) and *spry4* (B1-647) expression, while another was co-stained for *ets2* (B1-647) and *ier2a* (B2-546) (Molecular Instruments). For *LOV* HCR-FISH experiments, the B1-647 amplifier was used.

### Confocal imaging

Embryos that underwent HCR-IF or HCR-FISH were transferred from PBS to 1.2% low melting agarose (LMA) in embryo medium using a glass pipette pump and mounted on a plastic Petri dish. Blastula and gastrula stage embryos were mounted animal pole up, and 24 hpf embryos were mounted laterally. Bud-stage embryos were mounted in random positions due to orientation ambiguity. After LMA solidified, samples were covered in PBS and imaged using an upright Zeiss LSM 800, with an W N-Archoplan 10x/0.3 objective at 0.7x zoom, using Zeiss Zen software. For all experiments, images were captured in bidirectional scanning mode with 16 bit-depth and a resolution of 512 x 512 pixels. Pixel dwell time and line averaging were held constant between all experiments. Between 60-150 z-slices were obtained per embryo, depending on the developmental stage and size, with 9 μm between slices. ppERK signal (detected with B5-647) was imaged with a 640 nm laser, DAPI with a 405 nm laser, and GFP with a 488 nm laser. FGF target genes stained with B2-546 amplifier (*dusp6* and *ier2a*) were imaged with a 561 nm laser, and embryos stained with B1-647 amplifier (*LOV*, *spry4*, and *ets2*) were imaged with a 640 nm laser. Laser power and digital gain were adjusted to optimize dynamic range for each imaging run. Each set of exposed embryos and their dark controls were always mounted on the same dish and imaged with the same imaging conditions. For 24 hpf embryos, tiling of 3-6 frames was used to acquire whole-embryo images.

### Data analysis and statistics

#### Visualization and HCR-IF/FISH intensity quantification pipeline

HCR-IF/FISH z-stacks were processed using a custom MATLAB-based pipeline described in (Iannucci et al., 2026) to quantify average ppERK or gene expression intensities and generate maximum intensity projections (MIPs) for display. The MATLAB code can be accessed here: https://www.nichd.nih.gov/research/atNICHD/Investigators/rogers/software. Average ppERK intensities were measured within the entire embryo, rather than just the nuclei, for two reasons: 1) we observed both cytoplasmic and nuclear ppERK signal, and 2) we were unable to cleanly segment nuclei from 24 hpf embryo images, and wished to apply an identical analysis approach to all datasets assessed here. For HCR-FISH experiments, average intensities were also measured within the entire embryo.

To visualize imaging data, the pipeline accepts user-defined minima and maxima for the brightness in each channel and generates MIPs with identical display conditions.

To quantify whole-embryo intensity in all embryos except 24 hpf, the pipeline 1) first generates a nuclear mask using user-defined thresholds in the DAPI channel, 2) applies a user-defined “dilation factor” to the nuclear mask to expand the nuclei to overlap, creating a new mask that encompasses the entire embryo (nuclei and cytoplasm) but excludes most of the pixels outside of the embryo, and 3) applies the whole-embryo mask to the relevant channel and measures the intensity of all pixels within this mask, and also calculates the average of these values. For 24 hpf embryos, we were unable to generate accurate embryo masks with this automated approach, we speculate due to differences in DAPI distribution and smaller nuclei. We therefore used manually-drawn ROIs to create masks for whole-embryo measurements from 24 hpf images. The average intensities directly calculated from each image by this pipeline are shown in Fig. S2-6.

#### Quality control and experiment structure

Individual MIPs were visually assessed using pre-established criteria prior to being included in downstream analyses. Images were excluded if over 1/3 of the embryo was damaged or obscured. For all except 24 hpf embryos, samples were also removed from consideration if there was another embryo or other foreign material in frame (because ROIs were manually drawn for 24 hpf embryos, these artifacts could be avoided).

Any embryo that passed screening was considered a technical replicate for its given exposure paradigm (or control group). Each set of embryos within a given treatment group in one trial was considered a single biological replicate. Three biological replicates were generated for each condition for all experiments. Each biological replicate contained 7-8 technical replicates (i.e., 7-8 dark and 7-8 light; most biological replicates contained 8 embryos). The sole exception was *LOV* HCR-FISH replicate 1, which contained 4 embryos per condition (replicates 2 and 3 contained 8 per condition). Unexposed or wild-type controls were collected, incubated, processed for HCR-IF/FISH, and imaged with their respective treatment group(s).

#### Uniformity comparisons

To compare spatial uniformity in ppERK signal, masked ppERK MIPs obtained from the MATLAB pipeline detailed above were run through a second MATLAB pipeline that extracted an array of all of the embryo-spanning pixels greater than a threshold of 300 AU. A histogram was generated from the array to visualize pixel intensity distribution. Further, the interquartile range (IQR) was calculated as the difference between the 75^th^ and 25^th^ percentiles. The IQR was then divided by the thresholded mean-ppERK pixel intensity of that image to normalize for differences in absolute intensity between embryos, generating the mean-adjusted interquartile range (MA-IQR).

#### Statistical analyses

Statistical analyses were performed in Excel and JMP (18.2.1-19.1.2) and graph generation was performed in GraphPad Prism (10.1.0 (264)).

To account for overall intensity differences between biological replicates arising from variation in HCR-IF/FISH and imaging sessions (e.g., see raw average intensities in Fig. S2-6), we calculated ppERK and gene-expression fold changes for each biological replicate. For each set of exposed embryos and their unexposed controls (or for *LOV* HCR-FISH experiments, heterozygous transgenics and wild-type controls), we first determined the mean of the average ppERK values from the set of unexposed controls. To calculate fold changes, we then divided each exposed embryo’s average ppERK intensity by this unexposed average. The resulting fold changes were then averaged to represent the mean fold change from each biological replicate as shown in Fig. 2C, 2F, 3B, 4B, and Fig. S6C.

Fold-change values were then analyzed by linear mixed-effects models (LMMs) for the following experiments: Analysis 1 - intensity comparisons (Fig. 2C), Analysis 2 - mean-adjusted interquartile range of pixel-intensity counts (Fig. 2D), Analysis 3 - gene expression intensity comparisons (Fig. 2F), Analysis 4 - exposure length comparisons (Fig. 3), Analysis 5 - stage comparisons (Fig. 4), and Analysis 6 - *LOV* mRNA quantification (Fig. S6C). Each LMM uses Standard Least Squares for the fixed effects and Restricted Maximum Likelihood (REML) for the estimation method. For each analysis, the biological replicate was a random effect. Where warranted by a significant (*p* < 0.05) fixed effect, post-hoc comparisons were conducted. For each analysis, fixed effects, their significance results, and their post-hoc significance tests are listed in Table S1. P-values are reported for post-hoc comparisons in Tables S2-S7.

---

**Fig. S1:**
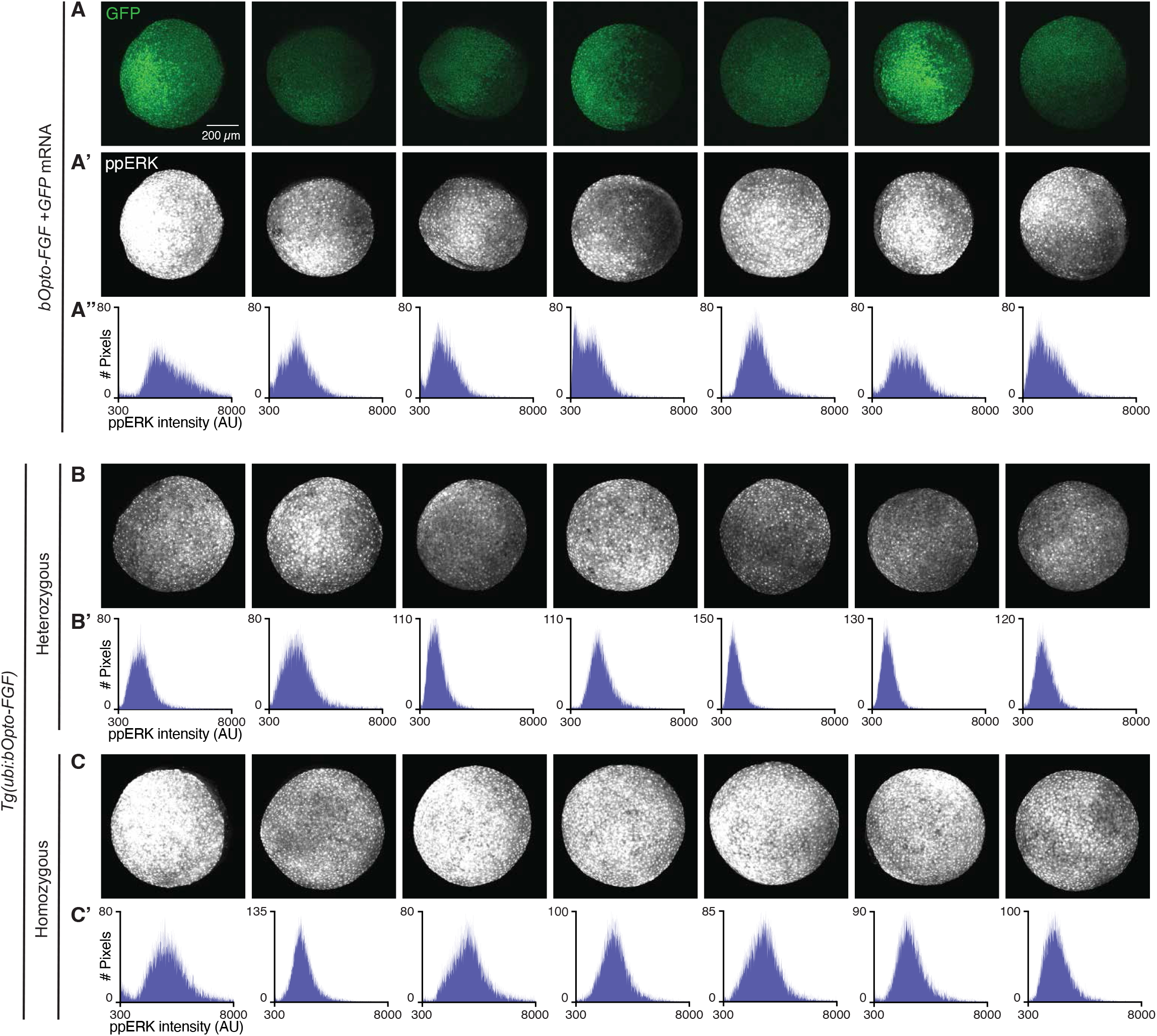
Spatial uniformity of optogenetic FGF/ERK signaling activation. *Tg(ubi:bOpto-FGF)* and *bOpto-FGF* + *GFP* mRNA-injected embryos were exposed to 455 nm light for 30 min beginning at shield stage. Embryos were fixed along with unexposed controls. HCR-IF staining for ppERK was used to quantify FGF/ERK signaling. A-A’) GFP signal (A) and ppERK signal (A’) from light exposed mRNA-injected embryos from one replicate. B,C) ppERK signal from light exposed heterozygous (B) and homozygous (C) embryos from one replicate. Scale bar = 200 µm. A”,B’,C’) Histograms of ppERK pixel intensities from above images. Note that y-axes may differ for display purposes.

**Fig. S2:**
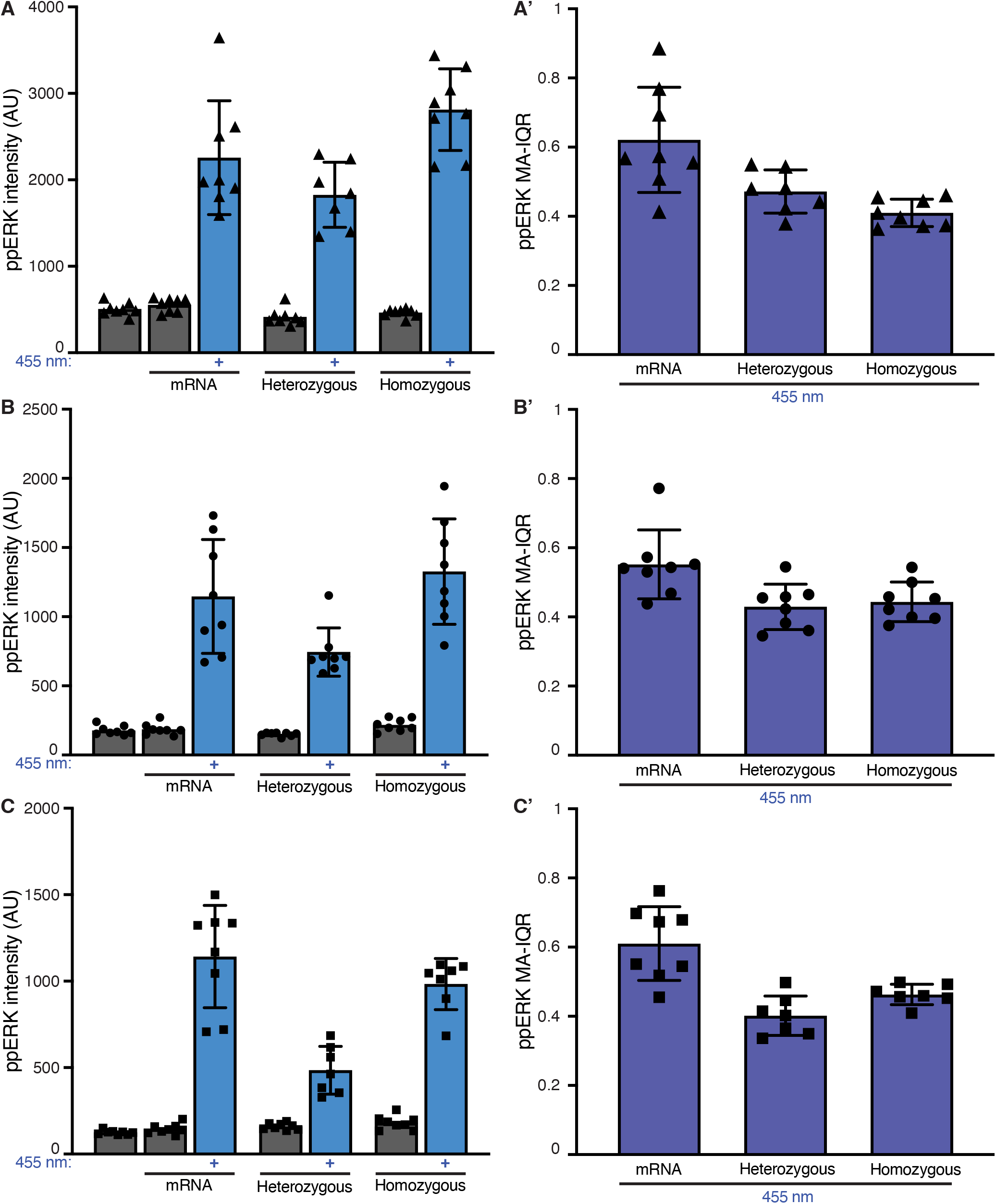
Optogenetic FGF/ERK activation replicates. *Tg(ubi:bOpto-FGF)* and *bOpto-FGF* + *GFP* mRNA-injected embryos were exposed to 455 nm light for 30 min beginning at shield stage. Embryos were fixed along with unexposed controls. mRNA replicates also included uninjected, unexposed controls. HCR-IF staining for ppERK was used to quantify FGF/ERK signaling. A,B,C) Raw ppERK intensities from three replicates. Each data point shows the average ppERK intensity from one embryo. A’,B’,C’) Corresponding ppERK mean-adjusted interquartile ranges (MA-IQR) from the three replicates shown in A, B, and C. Each data point shows the MA-IQR from one embryo. For all graphs, the symbol used for each replicate is consistent with Fig. 2, and error bars show SD.

**Fig. S3.**
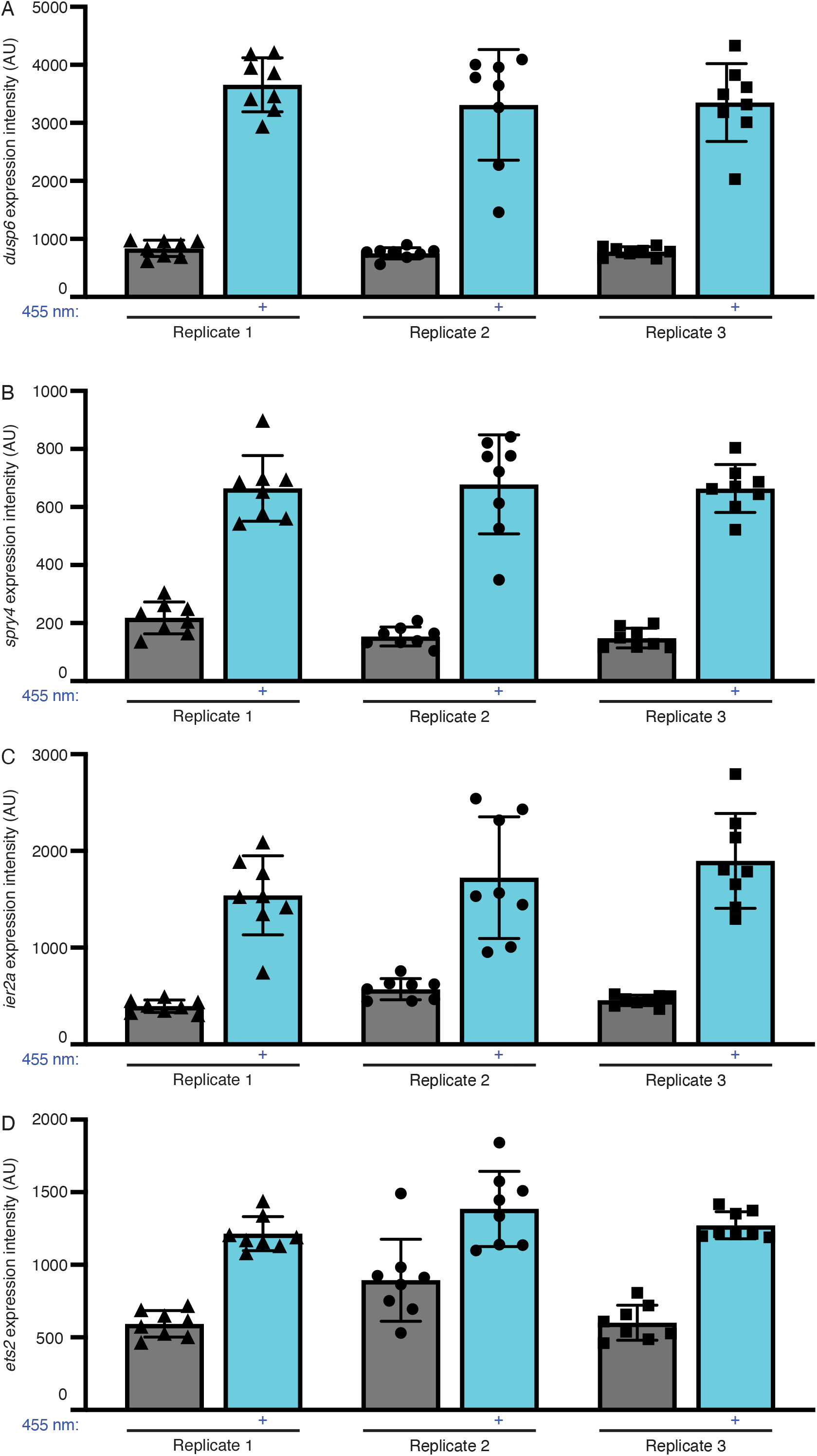
Light-mediated activation of FGF target gene expression. *Tg(ubi:bOpto-FGF)* heterozygous embryos were exposed to 455 nm light for 45 min beginning at 50% epiboly. Exposed embryos were fixed along with unexposed controls and HCR-FISH was performed for the indicated genes. A-D) Raw HCR-FISH intensity data for each gene from three replicates. Each data point shows the average intensity from one embryo. The symbol used for each replicate is consistent with Fig. 2. Error bars show SD.

**Fig. S4:**
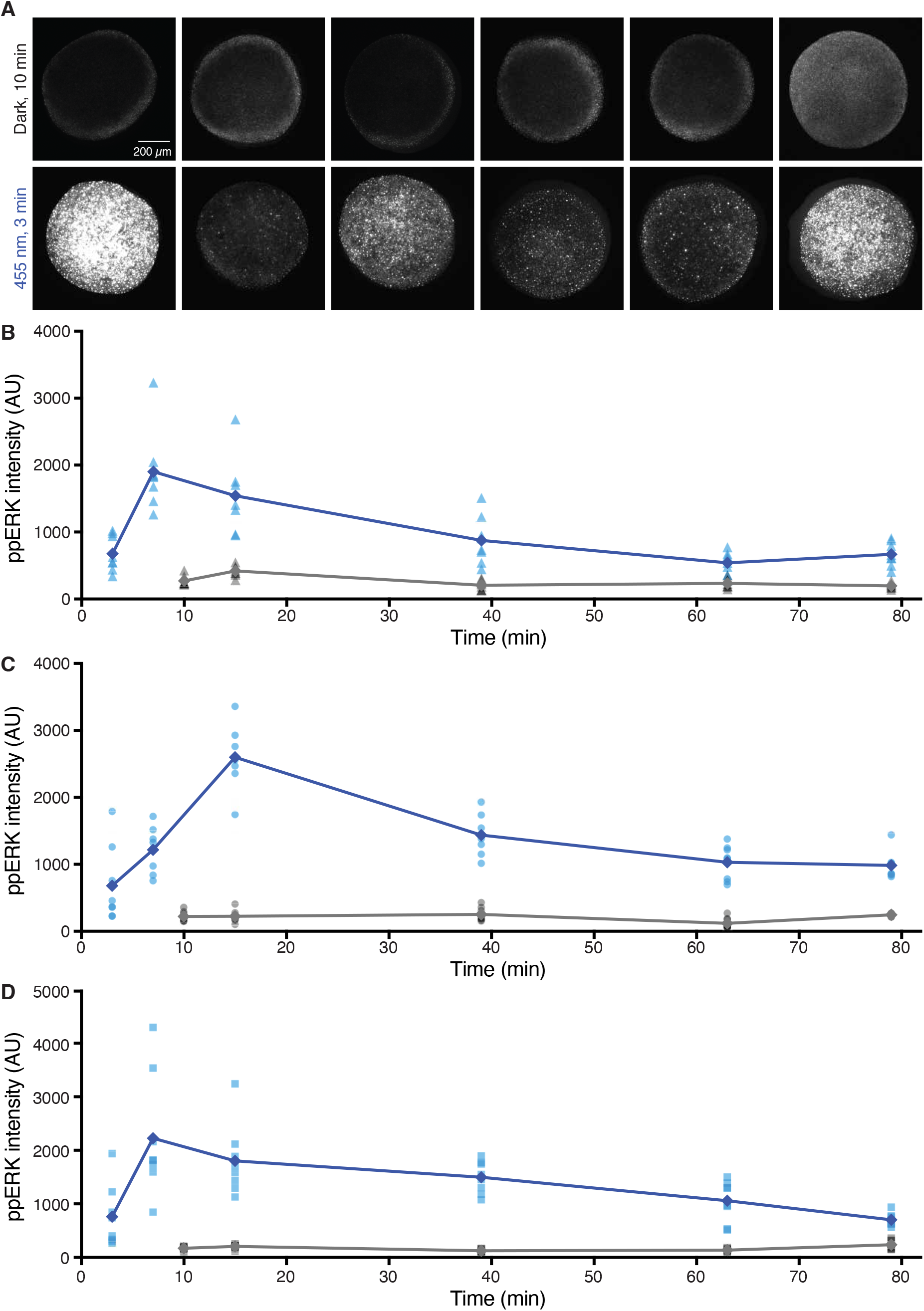
Kinetics independent replicates & representative 3 min images. Heterozygous *Tg(ubi:bOpto-FGF)* embryos were exposed to 455 nm light beginning at 30% epiboly. Embryos were fixed along with time-matched unexposed controls at the indicated durations. HCR-IF ppERK staining was used to assess FGF/ERK activity. A) Representative images of embryos exposed to blue light and fixed at 3 min (bottom row) and dark controls fixed at 10 min (top row). Scale bar = 200 µm. B-D) Raw ppERK intensity data from three replicates. Each data point (triangle, circle, or square) shows the average ppERK intensity from one embryo. The symbol used for each replicate is consistent with Fig. 3. Diamonds represent averaged data.

**Fig. S5:**
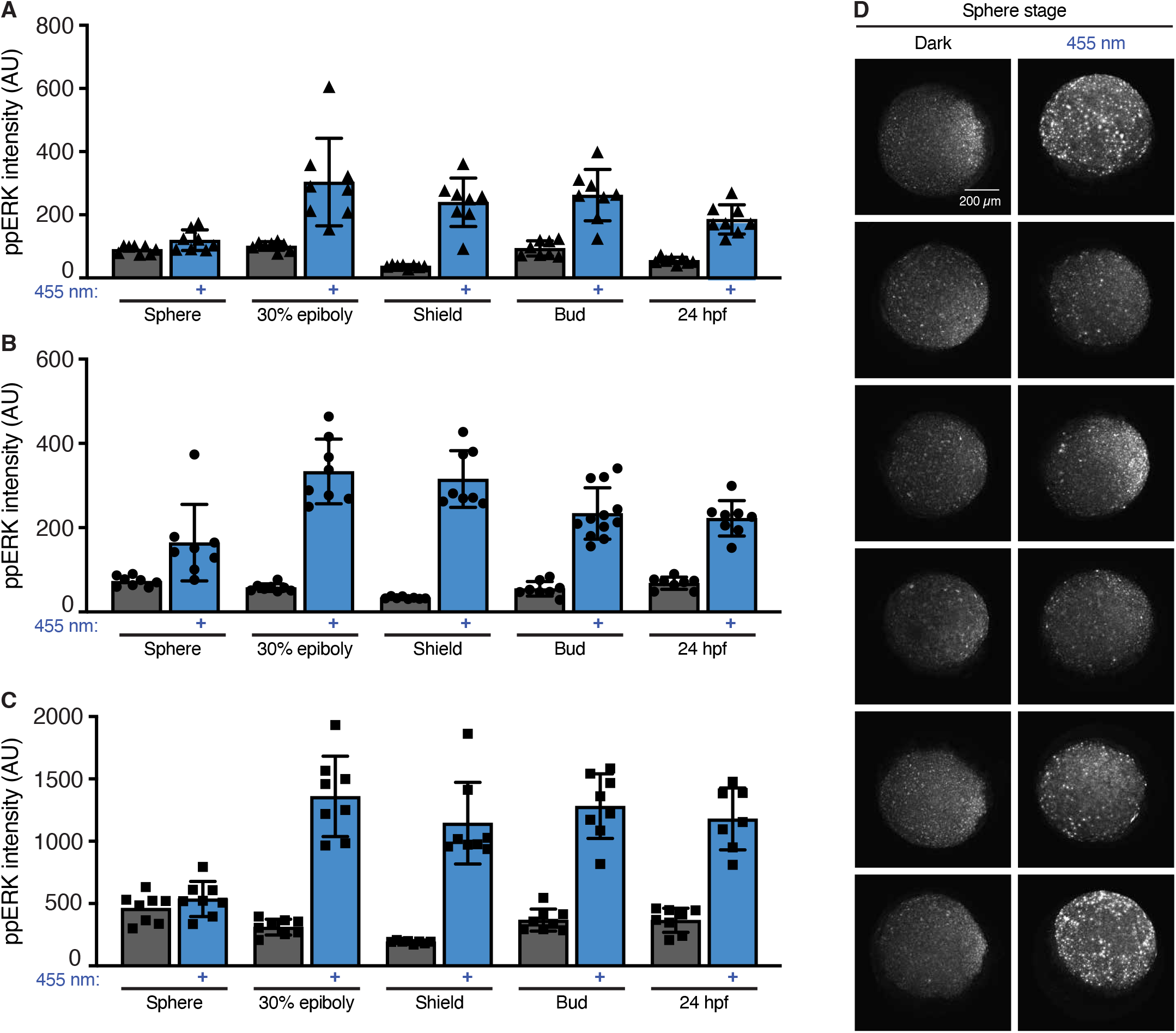
Timecourse replicates & representative sphere stage images. Heterozygous *Tg(ubi:bOpto-FGF)* embryos were exposed to 455 nm light beginning at the indicated stages. Embryos were fixed after 30 min along with stage-matched unexposed controls. HCR-IF for ppERK was used to detect FGF/ERK signaling activity. A-C) Raw ppERK intensity data from three replicates. Each data point shows the average ppERK intensity from one embryo. The symbol used for each replicate is consistent with Fig. 4. Error bars show SD. D) Representative images of blue light-exposed sphere-stage heterozygous embryos (right) and unexposed controls (left). Scale bar = 200 µm.

**Fig. S6.**
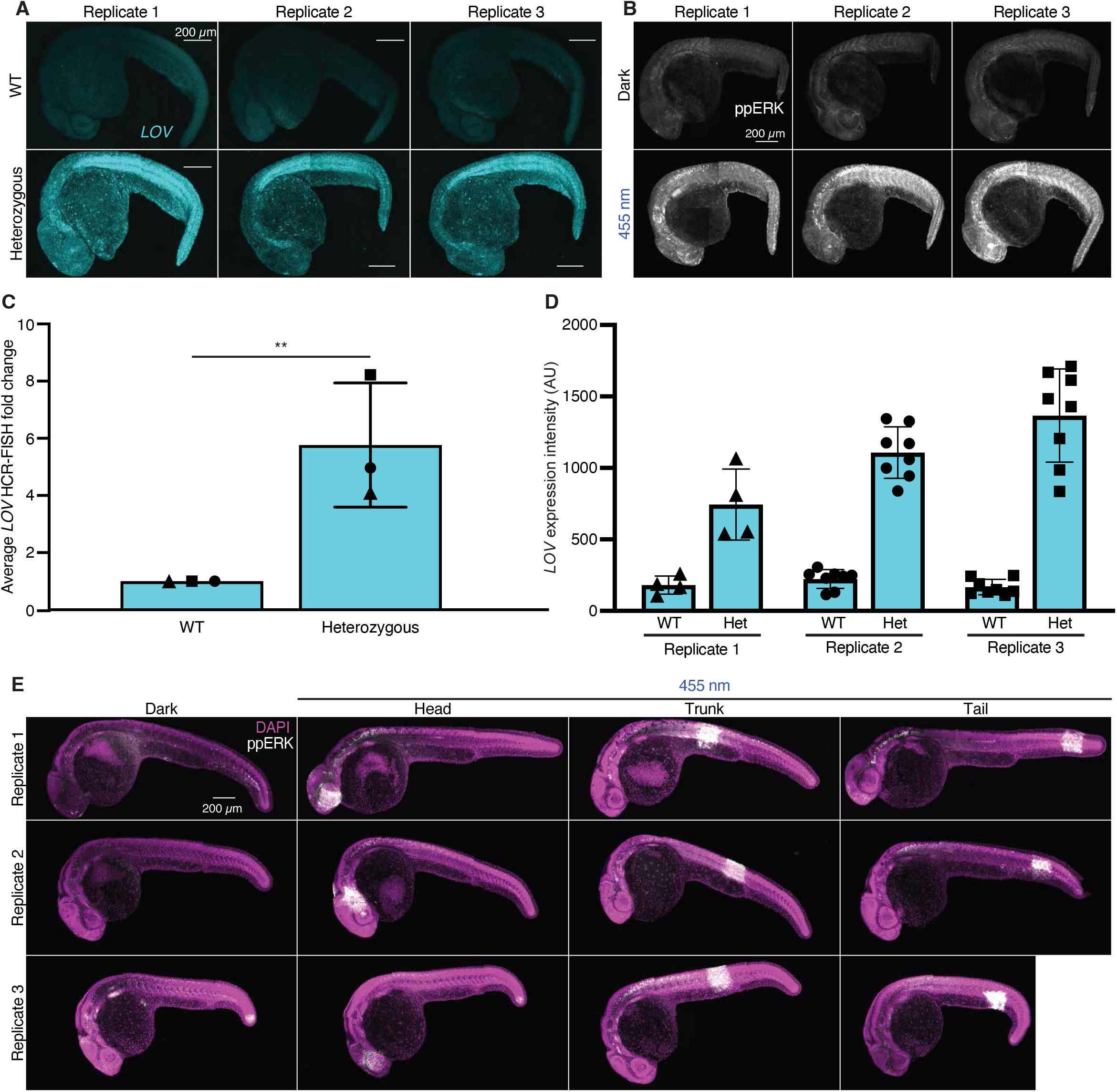
Distribution of bOpto-FGF mRNA & FGF/ERK activation after uniform or localized light exposure at 1 dpf. A, C and D: Dark-reared wild-type (top row) and *Tg(ubi:bOpto-FGF)* heterozygous embryos (bottom row) were fixed at 24 hpf. HCR-FISH was used to detect *bOpto-FGF* mRNA using probes targeting the *LOV* domain. A) Representative images from three replicates are shown. B) *Tg(ubi:bOpto-FGF)* heterozygous embryos were raised in the dark until 24 hpf, then either exposed to 455 nm light for 30 min (bottom row) or kept in the dark (top row), then stained for ppERK. Representative images from three replicates shown. C) Average intensity fold changes for three replicates of HCR-FISH for *LOV* mRNA. D) Raw intensity data from three *LOV* HCR-FISH replicates. Each data point shows the average intensity from one embryo. The symbol used for each replicate is consistent with C. Error bars show SD. E) *Tg(ubi:bOpto-FGF)* heterozygous embryos were raised in the dark until 1 dpf. Embryos were then exposed to a 170 x 170 µm square of 455 nm light for 1 min using a digital micromirror device mounted on a confocal microscope. Representative merged images showing DAPI (magenta) and ppERK (gray) staining from three independent replicates are shown for head-, trunk-, or tail-exposed embryos, and dark controls. Scale bar = 200 µm.

**Table S1:** List of fixed effects and post-hoc analyses used.

| Analysis | Figure | Fixed effects + p-values | Post-hoc analysis |
| --- | --- | --- | --- |
| 1 – ppERK intensity comparisons | 2C | 1. Exposure (dark or blue) ($p < 0.0001$)<br>2. Embryo type (Injected, het, or hom) ($p = .00036$)<br>3. Exposure*embryo type ($p = .00036$) | Tukey's Honest Significant Difference (HSD) |
| 2 – Mean-adjusted interquartile range | 2D | 1. Embryo type ($p < 0.0001$) | Tukey's HSD |
| 3 – Gene expression intensity comparisons | 2F | 1. Exposure (dark or blue) ($p < 0.0001$ for both co-stained sets)<br>2. Gene ($p = 0.037$ for *dusp6/spry4* and $p < 0.0001$ for *ets2* and *ier2a*)<br>3. Exposure*gene ($p = 0.037$ for *dusp6/spry4* and $p < 0.0001$ for *ets2* and *ier2a*) | Tukey's HSD |
| 4 – Duration comparisons | 3B | 1. Exposure ($p < 0.0001$)<br>2. Exposure duration ($p < 0.0001$)<br>3. Exposure*exposure duration ($p < 0.0001$) | Student's t-test with a Bonferroni correction |
| 5 – Stage comparisons | 4B | 1. Exposure ($p < 0.0001$)<br>2. Stage ($p < 0.0001$)<br>3. Exposure*Stage ($p < 0.0001$) | Tukey's HSD |
| 6 – LOV mRNA quantification | S6D | 1. WT vs. Het ($p = 0.0078$) | N/A (only one comparison made) |

**Table S2:**
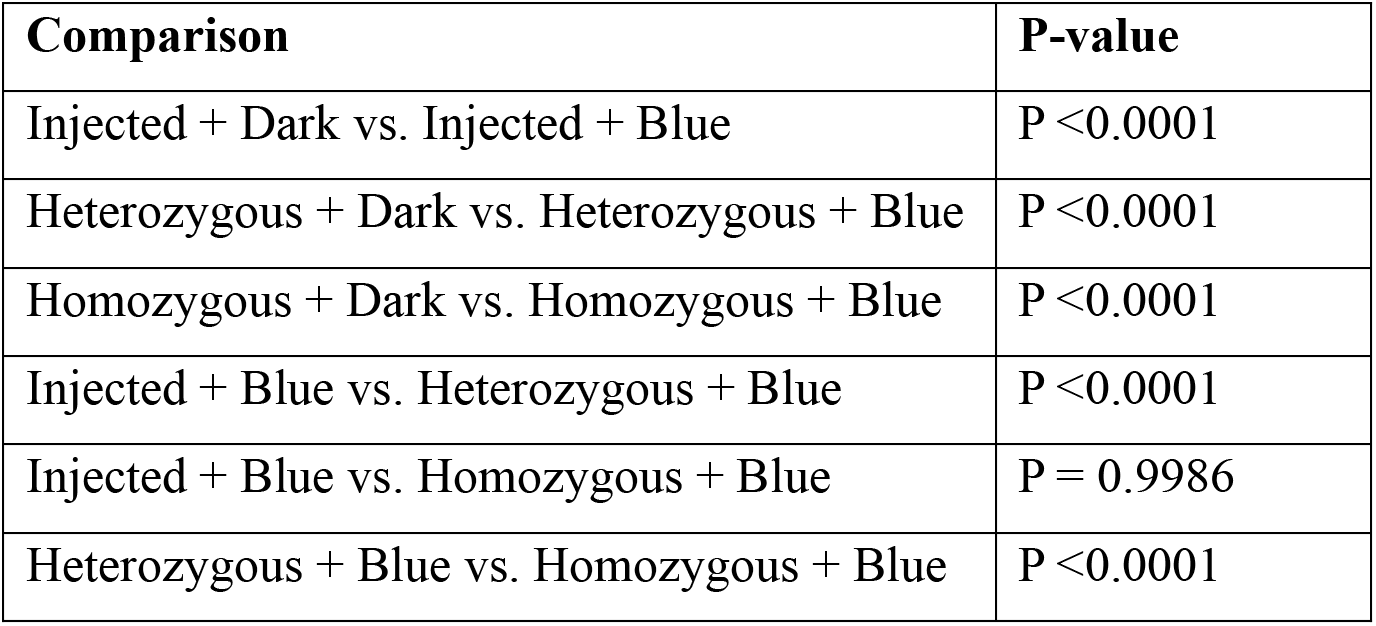
Corrected P-values from intensity analysis in Fig. 2C. (Analysis 1: Linear mixed-effects model with Tukey’s HSD to correct for multiple comparisons)

| Comparison | P-value |
| --- | --- |
| Injected + Dark vs. Injected + Blue | $P < 0.0001$ |
| Heterozygous + Dark vs. Heterozygous + Blue | $P < 0.0001$ |
| Homozygous + Dark vs. Homozygous + Blue | $P < 0.0001$ |
| Injected + Blue vs. Heterozygous + Blue | $P < 0.0001$ |
| Injected + Blue vs. Homozygous + Blue | $P = 0.9986$ |
| Heterozygous + Blue vs. Homozygous + Blue | $P < 0.0001$ |

**Table S3:** Corrected P-values from mean-adjusted interquartile range analysis in Fig. 2D. (Analysis 2: Linear mixed-effects model with Tukey’s HSD to correct for multiple comparisons)

| Comparison | P-value |
| --- | --- |
| Injected + Blue vs. Heterozygous + Blue | P <0.0001 |
| Injected + Blue vs. Homozygous + Blue | P <0.0001 |
| Heterozygous + Blue vs. Homozygous + Blue | P = 0.9852 |

**Table S4:** Corrected P-values from HCR-FISH in Fig. 2F. (Analysis 3: Linear mixed-effects model with Tukey’s HSD to correct for multiple comparisons)

| Comparison | P-value |
| --- | --- |
| *dusp6*: Dark vs. Blue | P <0.0001 |
| *spry4*: Dark vs. Blue | P <0.0001 |
| *ets2*: Dark vs. Blue | P <0.0001 |
| *ier2a*: Dark vs. Blue | P <0.0001 |

**Table S5A:** Dark vs. blue corrected p-values for kinetics analysis in Fig. 3B. (Analysis 4: Linear mixed-effects model with Bonferroni corrections following Student’s t-tests)

| Comparison | P-value |
| --- | --- |
| 10 min + Dark vs. 3 min + Blue | P = 0.0155 |
| 10 min + Dark vs. 7 min + Blue | P <0.0001 |
| 15 min + Dark vs. 15 min + Blue | P <0.0001 |
| 39 min + Dark vs. 39 min + Blue | P <0.0001 |
| 63 min + Dark vs. 63 min + Blue | P <0.0001 |
| 79 min + Dark vs. 79 min + Blue | P = 0.0081 |

**Table S5B:** Time point comparison corrected p-values for analysis in Fig. 3B. (Analysis 4: Linear mixed-effects model with Bonferroni corrections following Student’s t-tests)

| **Comparison** | **7 min** | **15 min** | **39 min** | **63 min** | **79 min** |
| --- | --- | --- | --- | --- | --- |
| 3 min | P <0.0001 | P <0.0001 | P <0.0001 | P = 0.0006 | P = 1.0 |
| 7 min | - | P = 1.0 | P = 1.0 | P = 0.0231 | P <0.0001 |
| 15 min | - | - | P = 1.0 | P = 0.1050 | P <0.0001 |
| 39 min | - | - | - | P = 1.0 | P <0.0001 |
| 63 min | - | - | - | - | P = 0.0013 |

**Table S6A:**
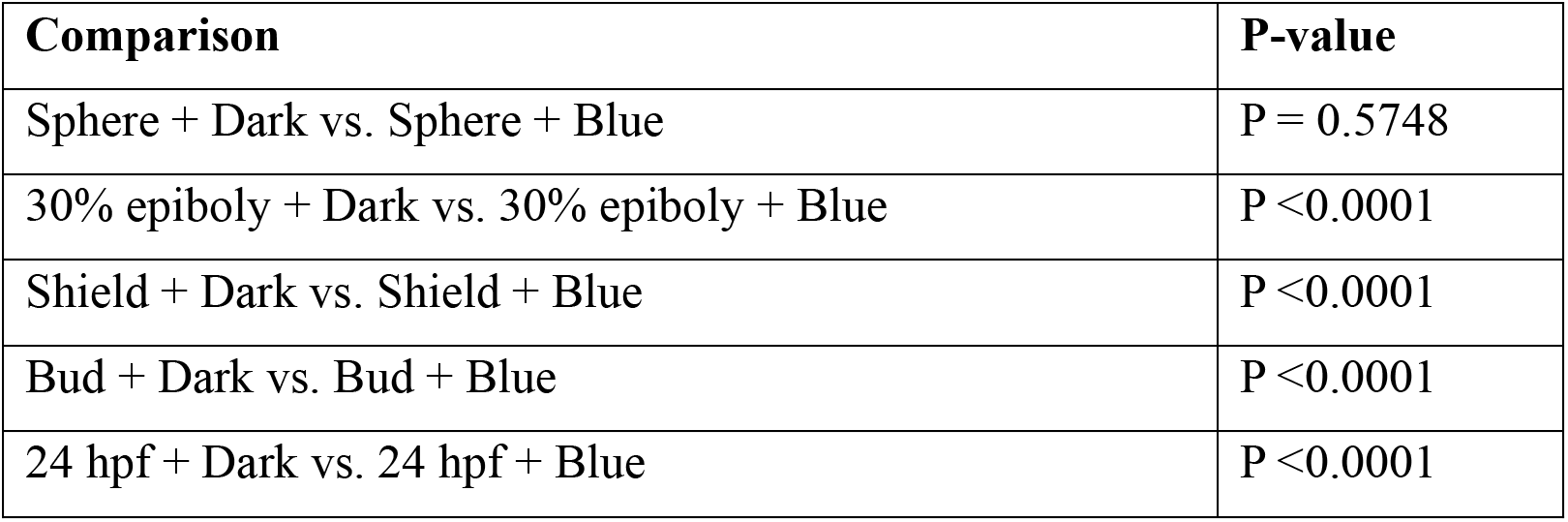
Dark vs. blue corrected p-values for stage analysis in Fig. 4B. (Analysis 5: Linear mixed-effects model with Tukey’s HSD to correct for multiple comparisons)

| **Comparison** | **P-value** |
| --- | --- |
| Sphere + Dark vs. Sphere + Blue | P = 0.5748 |
| 30% epiboly + Dark vs. 30% epiboly + Blue | P <0.0001 |
| Shield + Dark vs. Shield + Blue | P <0.0001 |
| Bud + Dark vs. Bud + Blue | P <0.0001 |
| 24 hpf + Dark vs. 24 hpf + Blue | P <0.0001 |

**Table S6B:** Stage comparison corrected p-values for analysis in Fig. 4B. (Analysis 5: Linear mixed-effects model with Tukey’s HSD to correct for multiple comparisons)

| **Comparison** | **30% Epiboly** | **Shield** | **Bud** | **24hpf** |
| --- | --- | --- | --- | --- |
| Sphere | P <0.0001 | P <0.0001 | P <0.0001 | P <0.0001 |
| 30% Epiboly | - | P <0.0001 | P = 0.0332 | P = 0.0058 |
| Shield | - | - | P <0.0001 | P <0.0001 |
| Bud | - | - | - | P = 0.9999 |

